# MicroRNA arm switching regulated by uridylation

**DOI:** 10.1101/2020.04.06.027813

**Authors:** Haedong Kim, Jimi Kim, Sha Yu, Young-Yoon Lee, Junseong Park, Ran Joo Choi, Seon-Jin Yoon, Seok-Gu Kang, V. Narry Kim

## Abstract

Strand selection is a critical step in microRNA (miRNA) biogenesis. Although the dominant strand may alter depending on cellular contexts, the molecular mechanism and physiological significance of such alternative strand selection (or “arm switching”) remain elusive. Here we find mir-324 as one of the strongly regulated miRNAs by arm switching, and identify terminal uridylyl transferases TUT4 and TUT7 as the key regulators. Uridylation of pre-mir-324 by TUT4/7 re-positions DICER on the pre-miRNA and shifts the cleavage site. This alternative processing produces a duplex with a different terminus, from which the 3′ strand (3p) is selected instead of the 5′ strand (5p). In glioblastoma, the TUT4/7 and 3p levels are upregulated while the 5p level is reduced. Manipulation of the strand ratio is sufficient to impair glioblastoma cell proliferation. This study uncovers a role of uridylation as a molecular switch in alternative strand selection and implicates its therapeutic potential.

## INTRODUCTION

MicroRNA (miRNA) serves as a guide molecule in RNA silencing through sequence-specific pairing with the targets. Because the miRNA sequence is embedded in the duplex region of its precursor, the guide strand should be selected from the duplex correctly to ensure the functionality (Khvorova et al., 2003; Schwarz et al., 2003). It was initially assumed that only one strand is dominantly chosen as a mature miRNA. However, the strand ratios of some miRNAs alter depending on the tissues, developmental stages, and pathological conditions (Chen et al., 2018; Chiang et al., 2010; Ro et al., 2007; Tsai et al., 2016; Zhang et al., 2019). These observations implicated there may be an active mechanism that controls alternative strand selection. However, the molecular principle underlying such “arm switching” remains to be elucidated.

The miRNA duplex is generated by consecutive actions of two RNase III enzymes. The nuclear RNase III DROSHA initiates miRNA maturation by processing the primary transcript (pri-miRNA). Together with its cofactor DGCR8, DROSHA forms a complex known as the Microprocessor that recognizes multiple sequence and structural motifs of pri-miRNA for precise cleavage (Denli et al., 2004; Gregory et al., 2004; Han et al., 2004; Lee et al., 2003). DROSHA introduces a staggered cut, liberating a small hairpin of ∼65 nt with a 2-nt 3′ overhang (called pre-miRNA). Following pre-miRNA export to the cytoplasm, DICER recognizes the 2-nt 3′ overhang on pre-miRNA using its platform and PAZ domains (Liu et al., 2018; Macrae et al., 2006; Tian et al., 2014). DICER acts like a molecular ruler that measures 22 nt from either the 5′ or 3′ end to determine the cleavage site on pre-miRNA (MacRae et al., 2007; Macrae et al., 2006; Park et al., 2011; Zhang et al., 2002). Pre-miRNA with a relatively unstable 5′ end (e.g., mismatch, G–U, or A–U pair) would be readily captured by the 5′ pocket (in the platform domain) of DICER and the cleavage site is dictated by the distance from the 5′ end (“5′-counting rule”) (Park et al., 2011; Tian et al., 2014). On the other hand, pre-miRNA with a stable 5′ end (e.g., G–C pair) relies on the interaction between its 3′ end and the 3′ pocket (in the PAZ domain) of DICER and is cleaved at the site ∼22 nt away from its 3′ end (“3′-counting rule”) (MacRae et al., 2007; Macrae et al., 2006; Park et al., 2011). The cleavage product is a ∼22-nt duplex with 2-nt 3′ overhangs at both ends, which is subsequently loaded onto an Argonaute (AGO) protein with the help from HSC70/HSP90 chaperone machinery (Iwasaki et al., 2010; Naruse et al., 2018). One strand (“guide”) remains in the AGO to direct gene silencing while the other strand (“passenger”) is expelled and degraded (Khvorova et al., 2003; Schwarz et al., 2003). The “seed” sequence (2–7 nucleotides relative to the 5′ end) of the guide strand is critical for target recognition (Bartel, 2018).

Alternative maturation of miRNA (such as processing by DROSHA/DICER at alternative sites) can yield multiple isoforms (called isomiRs) from the same hairpin and adds another layer of complexity to the control of gene expression. IsomiRs with distinct 5′ ends (5′-isomiRs) are of particular importance as the change at the 5′ end shifts the seed region and, therefore, has a significant impact on target specificity (Chiang et al., 2010; Fukunaga et al., 2012; Tan et al., 2014). The 5′ end heterogeneity can also alter the terminal properties of miRNA duplex and thus influence strand selection (Lee and Doudna, 2012; Wilson et al., 2015; Wu et al., 2009).

Strand selection is mainly determined by two rules involving thermodynamic characteristics and 5′ nucleotide identity of a miRNA duplex. The crystal structures of human AGO and its homologues have highlighted a key role of a basic pocket of the MID domain that interacts with the mono-phosphorylated 5′ end of the guide strand (Elkayam et al., 2012; Ma et al., 2005; Parker et al., 2005; Schirle and MacRae, 2012). Thermodynamically unstable 5′ ends would be more readily accessible to the pocket, leading to the preference for unstable 5′ ends by AGO (Khvorova et al., 2003; Schwarz et al., 2003). Furthermore, analyses of miRNA sequences indicated that there is a clear bias for U or A at the 5′ terminal position (Ghildiyal et al., 2010; Lau et al., 2001). The human AGO MID domain preferentially interacts with UMP or AMP but discriminates against CMP and GMP (Frank et al., 2010). Therefore, a strand with low thermodynamic stability and/or A/U at the 5′ end is favorably selected by AGO (Suzuki et al., 2015).

Despite these two rules, alternative strand selection or “arm switching” seems to take place considerably for certain miRNAs in a context-dependent manner (Chen et al., 2018; Chiang et al., 2010; Ro et al., 2007). For instance, miR-155-5p and miR-155-3p have opposite effects on type I interferon production (Alivernini et al., 2017), and the strand ratio of miR-155 changes throughout the activation stages of dendritic cells (Zhou et al., 2010). Arm switching has also been suggested as one of the miRNA diversification mechanisms during evolution (de Wit et al., 2009; Griffiths-Jones et al., 2011). Arm switching leads to accumulation of mature miRNAs with completely different seed sequences and, as a consequence, drastically changes the target repertoire of the miRNA gene. Therefore, unraveling the molecular mechanism of arm switching would be important to understand miRNA biogenesis regulation and the regulatory roles of miRNAs. However, it remains to be investigated if arm switching is indeed an active process, and if so, how it is regulated mechanistically.

In this study, we aimed to identify miRNAs that undergo conserved arm switching and uncover its molecular mechanism. We find that miR-324 is the most prominently regulated miRNA through arm switching and that the arm switching is actively controlled by uridylation enzymes.

## RESULTS

### Arm switching of miR-324

In order to gain molecular insights into miRNA arm switching, we first searched for miRNAs that exhibit significant alterations in their strand ratio. For accurate quantification of strand ratio, we employed the recently optimized protocol called AQ-seq that minimizes ligation bias in small RNA-seq (sRNA-seq) library construction (Kim et al., 2019). The variation of strand selection was estimated across 15 different mouse tissues/developmental stages (Figure 1A, median absolute deviation of the fraction of the 5p strand for each given miRNA hairpin). We also used a sRNA-seq dataset from 9 human cell lines to examine strand usage in humans (Figure S1A) (Mayr and Bartel, 2009). In both human and mouse datasets, the majority of miRNAs are invariable in strand ratio (∼86%, median variance below 3%), indicating that not every miRNA is subjected to arm switching. Nevertheless, we identified several intriguing cases unambiguously displaying substantial variations, such as miR-324, miR-362, miR-193a, and miR-140.

**Figure 1.**
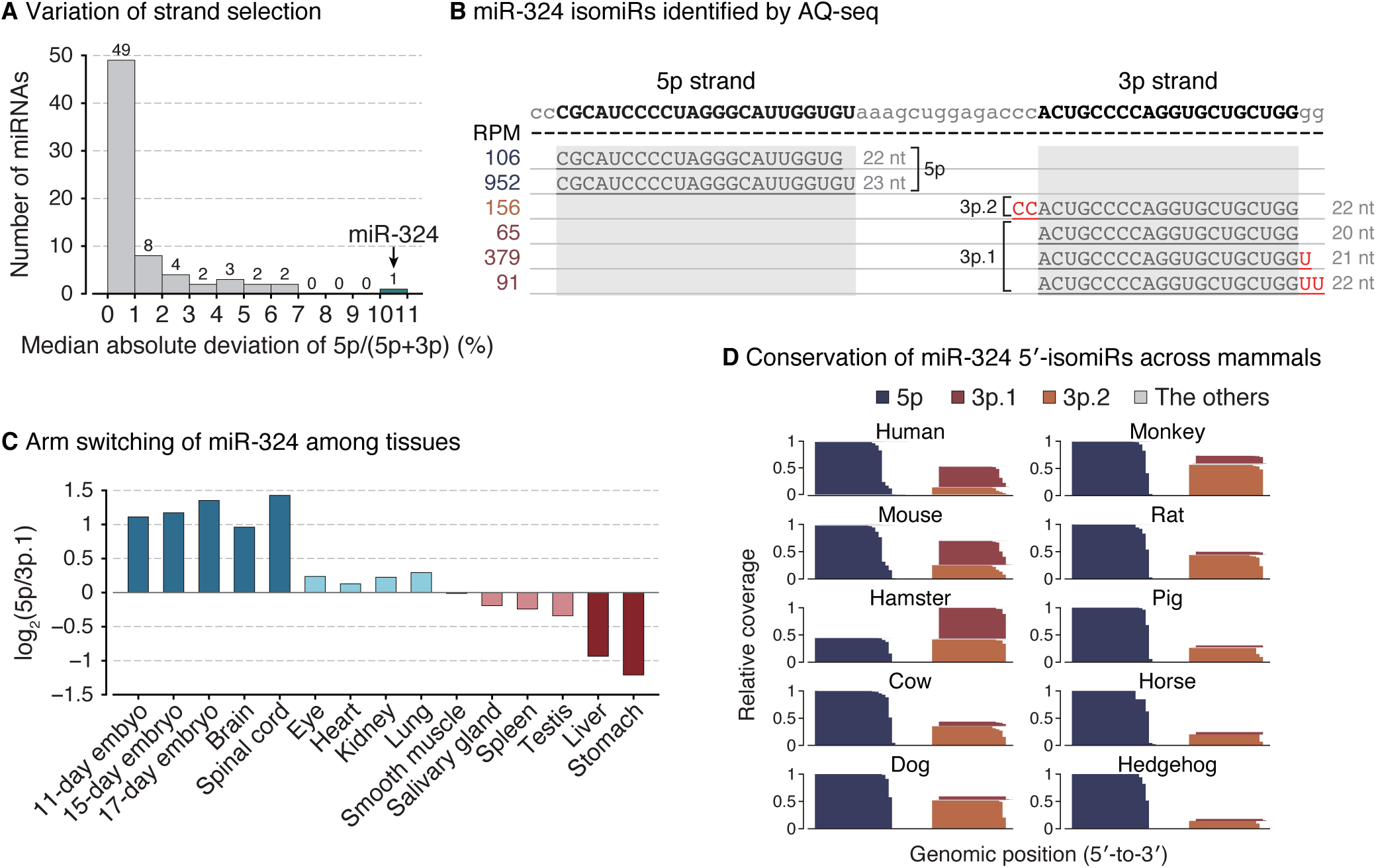
Arm switching of miR-324. **(A)** Median absolute deviation of the 5p proportion for a given miRNA in mouse tissues. Abundant and ubiquitous miRNAs (> 100 median RPM in both mouse tissue and human cell line datasets, > 0 RPM in all samples) were included in this analysis. **(B)** IsomiR profiles of miR-324 in HEK293T, uncovered by AQ-seq. RPM-normalized read counts are denoted on the left. Reference sequences of 5p and 3p are marked by grey shade. **(C)** Strand ratios (5p/3p.1) of miR-324 for the indicated panel of mouse tissues as measured by AQ-seq. **(D)** Conservation of the 5′-isomiRs of miR-324 in mammals. sRNA-seq reads of human, monkey, mouse, rat, hamster, cow, and horse were obtained from miRBase release 22 (Kozomara et al., 2019). For the other species, sRNA-seq reads were obtained from MirGeneDB release 2.0 (Fromm et al., 2019). RPM, reads per million. See also Figure S1.

miR-324 exhibits prominent arm switching in both human and mouse (Figures 1A–C and S1A–B). It was previously shown that miR-324-5p is downregulated in hepatocellular carcinoma and glioblastoma and suppresses cell proliferation and invasion (Cao et al., 2015; Zhi et al., 2017). In contrast, miR-324-3p is upregulated in hepatocellular carcinoma and promotes cell proliferation (Tuo et al., 2017; Xiao et al., 2017). These findings collectively suggest that two strands from the miR-324 hairpin may have distinct, possibly opposing, functions. Thus, the arm switching of miR-324 may have a substantial impact on cellular fate decision.

Notably, we detected three different groups of 5′-isomiRs (5p, 3p.1, and 3p.2) that are produced from the single MIR324 locus (Figure 1B). The 5p strand is dominant in mouse embryos and neuronal tissues while the major 3p isoform (3p.1) is prevalent in liver and stomach (Fig. 1C). The 5′-isomiRs have distinct seed sequences and, consequently, bind to different targets according to the miRNA-target interactome data (Figure S1C) (Helwak et al., 2013; Moore et al., 2015). Moreover, the 5′-isomiR groups are found across mammals, suggesting that the mechanism for alternative maturation is conserved evolutionarily (Figures 1D and S1D).

### miR-324 arm switching is controlled by TUT4 and TUT7

To understand the underlying mechanism for alternative strand usage, we examined sRNA-seq (AQ-seq) data because the reads may reflect the maturation processes of miRNAs, including 3′ end modifications and alternative processing events (Kim et al., 2019). Intriguingly, 3p.1 is highly uridylated (∼88% of 3p.1 reads) (Figure 1B). The uridylation frequency of 3p correlates negatively with the 5p/3p.1 ratio (Figure 2A). These observations led us to hypothesize that uridylation may be involved in the arm switching.

**Figure 2.**
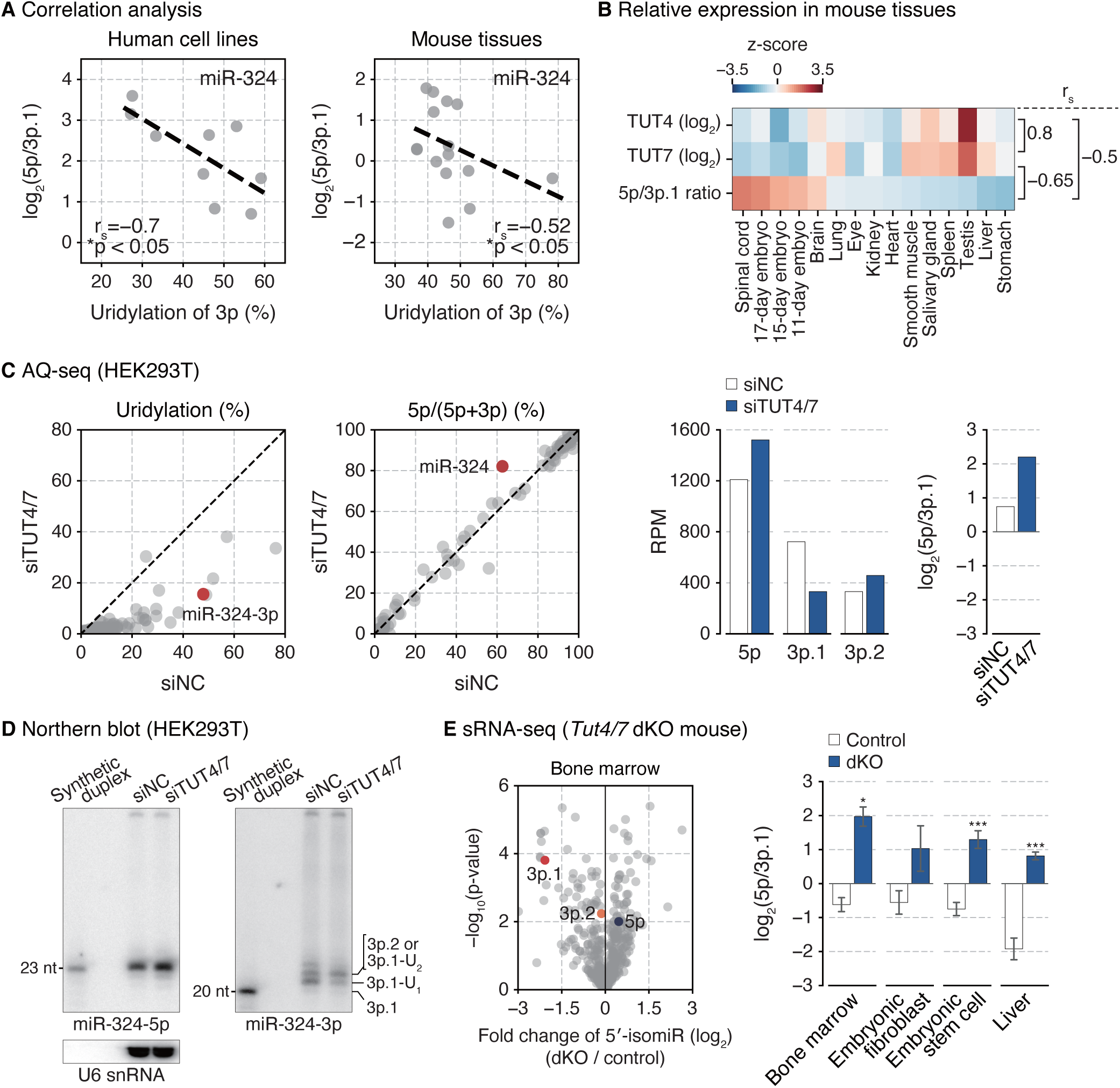
miR-324 arm switching is controlled by TUT4 and TUT7. **(A)** Negative association between the uridylation frequency of miR-324-3p and the 5p/3p.1 ratio of miR-324 in nine human cell lines and fifteen mouse tissues. The linear regression is shown with dashed lines. r_s_, Spearman correlation coefficient. *p < 0.05 by the two-sided test. **(B)** The relative TUT4/7 levels and the 5p/3p.1 ratios of miR-324 in the indicated panel of mouse tissues. The TUT4/7 mRNA levels and 5p/3p.1 ratios were quantified by RT-qPCR and AQ-seq, respectively. r_s_, Spearman correlation coefficient. **(C)** AQ-seq following knockdown of TUT4/7 in HEK293T. First and second panels: Scatter plots of uridylation frequency and the 5p proportion of miRNAs. Abundant miRNAs (> 100 RPM) in the siNC-transfected sample were included in the analysis. Third and fourth panels: Relative abundance of three 5′-isomiRs and log_2_-transformed 5p/3p.1 ratio of miR-324. **(D)** Northern blot of miR-324-5p and miR-324-3p in HEK293T. Synthetic miR-324 duplex composed of 23 nt-long 5p and 20 nt-long 3p.1 was used for size references. U6 snRNA was detected as a loading control. The same blot was probed for all the results. **(E)** sRNA-seq following double knockout of *Tut4/7* in mice. Left: A volcano plot showing abundance changes of 5′-isomiRs by TUT4/7 depletion in bone marrow. 5′-isomiRs with RPM over 10 in the control samples were included in this analysis. P-value by the two-sided Student’s *t* test. Right: log_2_-transformed 5p/3p.1 ratio of miR-324. Bars indicate mean ± s.d.. Bone marrow, n = 2; embryonic fibroblast, n = 2; embryonic stem cell, n = 3; control and *Tut4/7* dKO in liver, n = 4 and 3, respectively. *p < 0.05, ***p < 0.001 by the two-sided Student’s *t* test. RPM, reads per million. See also Figure S2.

Terminal uridylyl transferases TUT4 (also known as ZCCHC11 and TENT3A) and TUT7 (also known as ZCCHC6 and TENT3B) catalyze uridylation of diverse RNA species including a specific set of pre-miRNAs (e.g., *let-7* precursors) (Hagan et al., 2009; Heo et al., 2012; Heo et al., 2009; Liu et al., 2014; Thornton et al., 2012). TUT4 and TUT7 (TUT4/7) act redundantly on most substrates (Heo et al., 2012; Labno et al., 2016; Le Pen et al., 2018; Lim et al., 2014; Pirouz et al., 2019; Thornton et al., 2012; Warkocki et al., 2018). We quantified TUT4/7 levels in mouse tissues by RT-qPCR and found that the 5p/3p.1 ratio is higher in cell types where the TUT4 and TUT7 levels are relatively low (Figure 2B). To test the involvement of TUT4/7 in miR-324 maturation, we depleted TUT4/7 in HEK293T cells and performed sRNA-seq (AQ-seq) (Kim et al., 2019). TUT4/7 knockdown reduced uridylation of miRNAs including miR-324-3p (Figure 2C, first panel). Importantly, TUT4/7 knockdown resulted in a change in isoform composition of miR-324, with a reduction of 3p.1 and an increase of 5p and the minor isoform 3p.2. Consequently, the ratio between the major isoforms (5p/3p.1) increased (Figure 2C, second–fourth panels). These sequencing data were consistent with the northern blot (Figures 2D and S2A) and RT-qPCR results (Figure S2B), which confirmed alterations in the abundance of miR-324 isoforms upon TUT4/7 knockdown.

Moreover, re-analysis of the published sRNA-seq data from the *Tut4/7* double knockout mice shows that the 3p.1 level decreases while 5p increases in the knockout (Figures 2E, left panel, and S2C) (Morgan et al., 2017). Consequently, a dramatic change in the strand ratio was observed in all tissues/cell types examined (bone marrow, embryonic fibroblasts, embryonic stem cells, and liver) (Figures 2E, right panel). Together, the results demonstrate that TUT4/7 actively regulate miR-324 isoform selection in favor of 3p.1.

Of note, we noticed that the residual 3p.1 reads in the knockout samples are mostly adenylated at the 3′ end (Figure S2D). This may be attributed to the activity of terminal adenylyltransferase TENT2 (also known as PAPD4, GLD2, and TUT2) that we have previously shown to play a complementing role in TUT4/7-depleted cells, assisting the processing of group II pre-*let-7* family (Heo et al., 2012).

### Uridylation leads to alternative DICER processing of pre-miR-324

We next sought to understand mechanistically how TUT4/7 modulate the miR-324 strand selection. TUT4/7 are known to modify pre-miRNAs rather than mature miRNAs (Chiang et al., 2010; Kim et al., 2019; Liu et al., 2014). Thus, we interrogated the effect of uridylation on pre-miRNA processing by performing *in vitro* DICER processing assays with synthetic pre-miR-324. Remarkably, when pre-miR-324 carries one or two extra uridine residues at the 3′ end, the DICER processing site is shifted by 3 nt (Figure 3A, from position A to position B). Unmodified pre-miR-324 mainly releases a longer duplex containing 5p and 3p.2 (Figure 3A, blue arrowheads) while uridylated pre-miR-324 is cleaved at the alternative position B, yielding a shorter duplex composed of shorter 5p and 3p.1 (Figure 3A, red arrowheads). Thus, TUT4/7-mediated uridylation of pre-miR-324 leads to alternative DICER processing. Consistent with this *in vitro* assay, the sequencing data from TUT4/7-depleted human and mouse cells showed that the relative abundance of the 3p isomiRs (3p.1 vs. 3p.2) changed upon TUT4/7 knockdown (Figures 2C, third panel, and 3B) and knockout (Figure 3C), corroborating the conclusion that uridylation alters DICER cleavage site selection.

**Figure 3.**
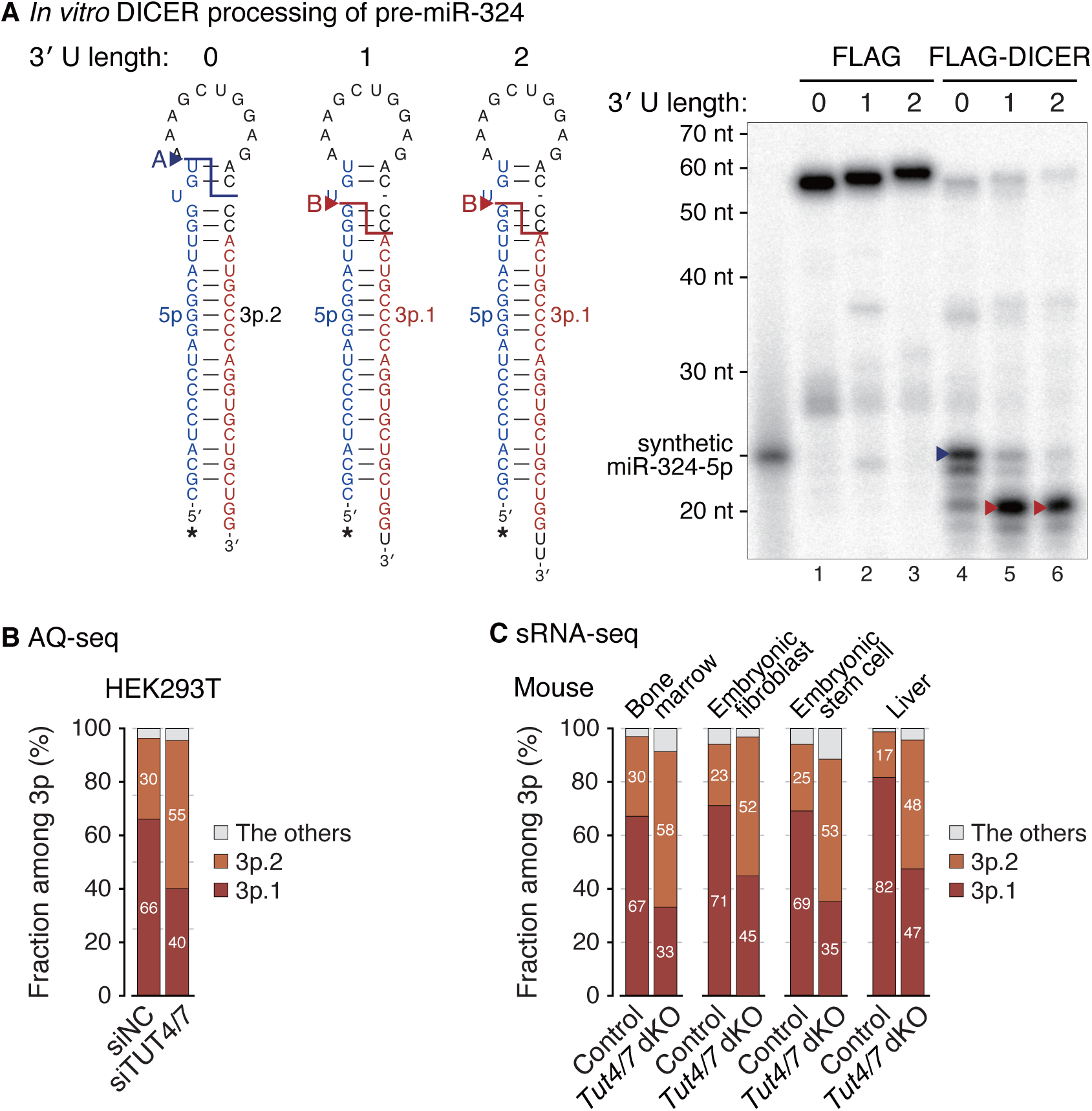
Uridylation leads to alternative DICER processing of pre-miR-324. **(A)** *In vitro* processing of unmodified, mono- or di-uridylated forms of pre-miR-324 by immunopurified DICER. Synthetic miR-324-5p (23 nt) marked in blue was used as a size control. Major cleavage products and their corresponding cleavage sites are marked with arrowheads. *, radiolabeled 5′ phosphates. **(B)** 5′-isomiR composition of miR-324-3p in HEK293T. **(C)** 5′-isomiR composition of miR-324-3p in mice. Bars indicate mean (bone marrow, n = 2; embryonic fibroblast, n = 2; embryonic stem cell, n = 3; control and *Tut4/7* dKO in liver, n = 4 and 3, respectively). See also Figure S3.

It was surprising to observe that uridylation of pre-miR-324 induced the 3-nt shift of the DICER processing site because pre-miR-324 has a terminal C–U mismatch and, hence, is predicted to follow the 5′-counting rule. According to the 5′-counting rule, cleavage site is dictated by the distance from the 5′ end and should not be affected by 3′ terminal uridylation (Figure S3) (Park et al., 2011). Therefore, this raised new questions concerning the DICER processing mechanism: (1) What causes the abrupt 3-nt shift in pre-miR-324 processing upon uridylation? (2) How do the established 5′- and 3′-counting rules apply to pre-miR-324 processing?

### Alternative DICER processing is facilitated by the double-stranded RNA binding domain (dsRBD)

We first investigated the determinant(s) responsible for the 3-nt shift. Structural modeling of human DICER in complex with pre-miR-324 suggested that the asymmetric U bulge near the cleavage site may cause steric hindrance against the RNase IIIb domain if the catalytic center of DICER is placed 1 nt away from the original cleavage site (Figure S4A) (Liu et al., 2018). To find out if the U bulge serves as an anti-determinant for cleavage at the neighboring position, we removed the bulge from pre-miR-324. For the “no-bulge” mutant, DICER no longer induced the abrupt shift to position B upon uridylation, but rather causing single-nucleotide gradual shifts (Figure 4A, lanes 10–12). Thus, the U bulge is an anti-determinant for cleavage at sites between positions A and B and thereby serves as the *cis*-acting element responsible for the unexpected 3-nt shift to position B.

**Figure 4.**
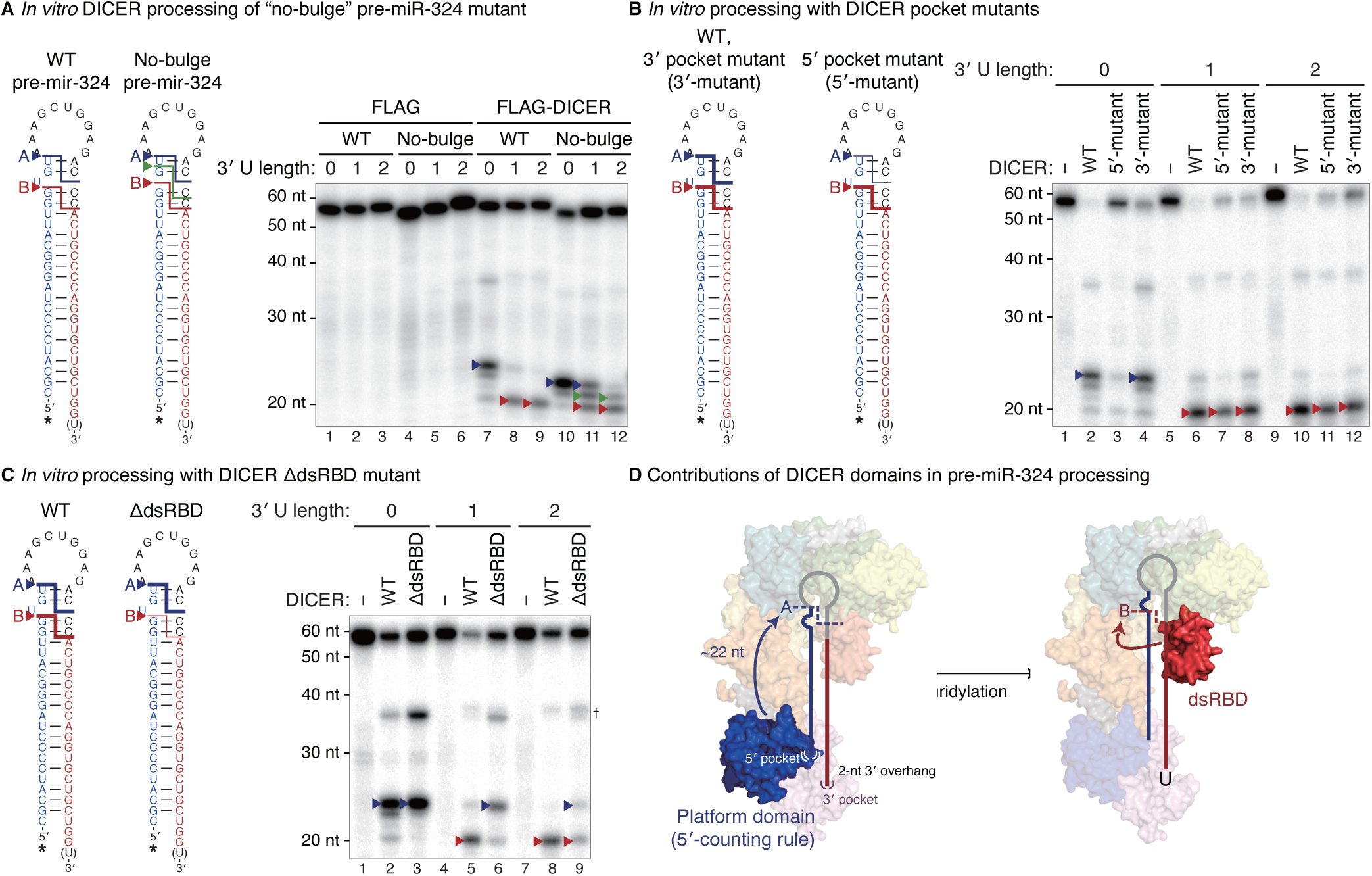
Alternative DICER processing is facilitated by the dsRBD. **(A)** *In vitro* processing of unmodified, mono- or di-uridylated forms of wild type or no-bulge mutant pre-miR-324 by immunopurified DICER. **(B–C)** *In vitro* processing of unmodified, mono- or di-uridylated forms of pre-miR-324 by immunopurified DICER pocket mutants (B) or dsRBD-deleted mutant (C). Major cleavage products and their corresponding cleavage sites are marked with arrowheads. *, radiolabeled 5′ phosphates. †, nicked products at the 3p positions. **(D)** A proposed model for the uridylation-mediated alternative DICER processing of pre-miR-324. See also Figure S4.

Next, to reveal how the 5′- and 3′-counting rules relate to pre-miR-324 processing, we utilized the DICER mutants in either the 5′ or 3′ pocket (Figure S4A) (Park et al., 2011). The 5′ pocket mutant DICER was severely impaired in cleaving pre-miR-324 at position A while the 3′ pocket mutant was only slightly affected in processing efficiency without an impact on cleavage site choice (Figure 4B). This result demonstrates the importance of the 5′ pocket for the cleavage at position A. Consistently, when the 5′ pocket mutant DICER was ectopically expressed in *Dicer*-null cells, it produced less miR-324-3p.2 (originating from position A) than the wild type DICER did, indicating that the 5′ pocket of DICER is required for cleavage at position A in cells (Figure S4B, blue arrowheads) (Park et al., 2011). In contrast, the cleavage at position B was not affected substantially by the 5′ pocket mutation (Figure S4B, red arrowheads). Thus, the cleavage at position A is indeed dictated by the 5′ pocket. However, if the 5′ pocket is the only determinant, how does 3′ uridylation lead to the shift to position B? Our results implicated that there must be a yet-unknown mechanism that contributes to DICER cleavage site selection.

This led us to investigate the contribution of other domains in DICER. We recently discovered that DROSHA uses the double-stranded RNA binding domain (dsRBD) to recognize a motif near the cleavage site to achieve precise processing (Fang and Bartel, 2015; Kwon et al., 2019). DROSHA has a similar overall structure to DICER, which suggests a common evolutionary origin of metazoan RNase III proteins (Kwon et al., 2016). To test the role of the DICER dsRBD *in vitro*, we generated a dsRBD-deletion mutant. Since the dsRBD is connected to the RNase IIIb domain via a flexible linker, the deletion is unlikely to alter the overall DICER conformation (Figure S4A). Intriguingly, the dsRBD deletion resulted in a significant change in the processing pattern (Figure 4C). Without the dsRBD, DICER cleaved mainly at position A, which indicates the dsRBD is responsible for the cleavage at position B. This result reveals a previously unappreciated role of the DICER dsRBD in cleavage site selection. Of note, the DICER-interacting partner, TRBP, has no impact on the uridylation-mediated alternative processing (Figure S4C, lanes 13–18).

Taken together, unmodified pre-miR-324 is cleaved by DICER at position A relying on the 5′ pocket in the platform domain (Figure 4D, left panel). However, when uridylated, the end structure (3–4 nt 3′ overhang) is unfit to be accommodated tightly by the platform/PAZ domains so that pre-miR-324 is repositioned in DICER with the help of the dsRBD, resulting in alternative processing at position B (Figure 4D, right panel).

### Alternative DICER processing leads to arm switching

Our results showed that pre-miR-324 can yield two alternative forms of duplexes: a long duplex (position A; 5p/3p.2) and a short duplex (position B; 5p/3p.1) (Figure 5A). Does this alternative processing lead to alternative strand selection? To examine strand selection in cells, we constructed reporters that contain the 3′ UTR complementary to either miR-324-5p or miR-324-3p.1 (Figure 5B, left panel). Synthetic duplexes (either the long duplex (5p/3p.2) or the short duplex (short 5p/3p.1)) were co-transfected with the reporters. The long duplex selectively repressed the reporter with the 5p complementary sites but not that with 3p.1 sites, confirming the production of 5p from the long duplex. Importantly, the short duplex downregulated the 3p.1 reporter but not the 5p reporter (Figure 5B, right panel). Accordingly, the long and short duplexes suppressed distinct predicted targets of 5p and 3p.1, respectively (Figures 5C and S5A–B) (Agarwal et al., 2015). These results show that 5p and 3p.1 are indeed differentially produced from the long duplex and the short duplex, respectively. Retrospective analysis of sRNA-seq data using the 5′ pocket mutant DICER also revealed that when the long duplex production is disturbed, the 5p/3p.1 ratio decreases (Figure S5C).

**Figure 5.**
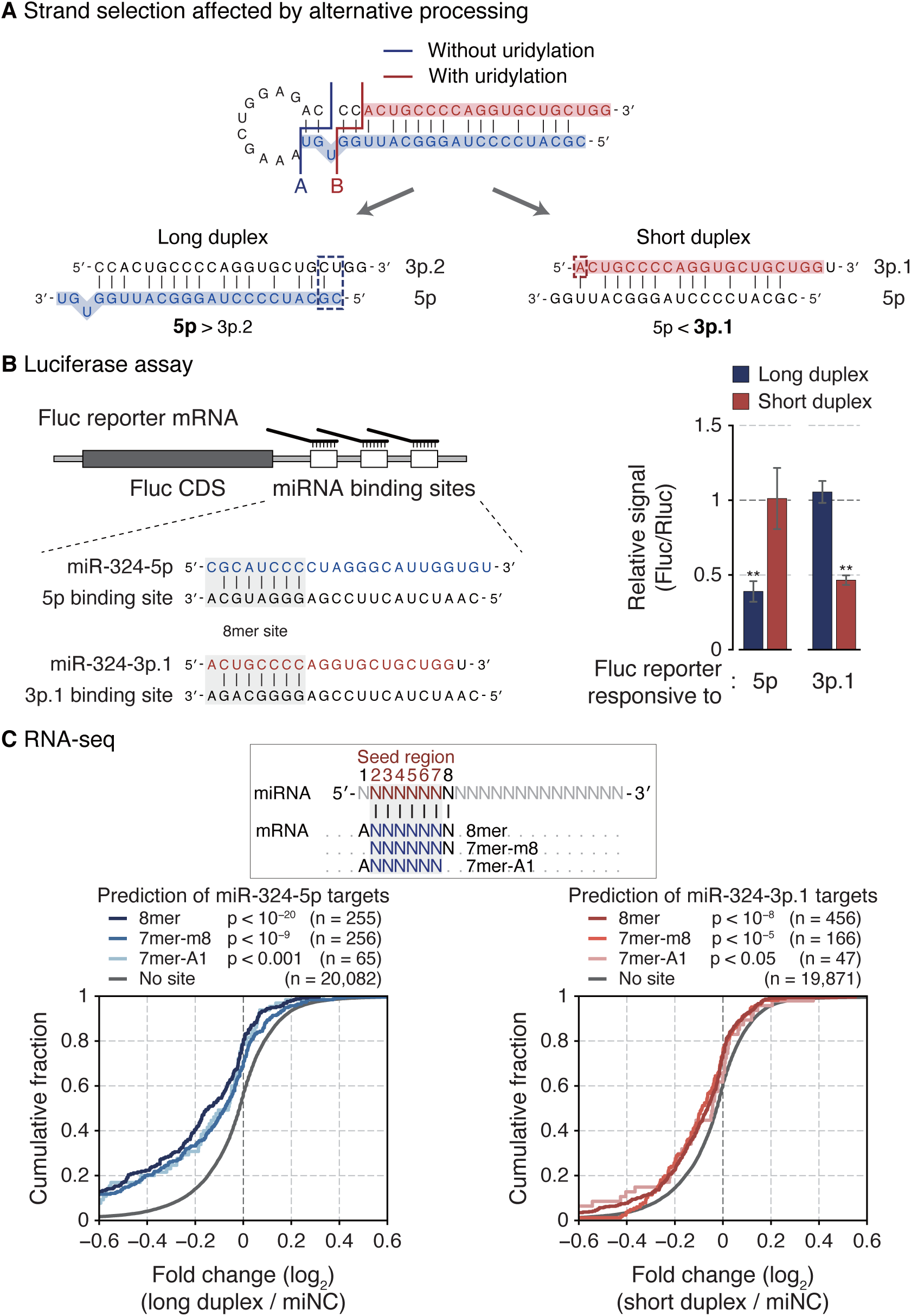
Alternative DICER processing leads to arm switching. **(A)** A schematic diagram of strand selection of miR-324 duplexes produced from unmodified and uridylated pre-miR-324. Dashed rectangles indicate end properties that dictate the strand selection of each duplex. **(B)** Luciferase reporter assay with two miR-324 duplexes. Left: An illustration of firefly reporter mRNAs that have three 8mer target sites of either miR-324-5p or 3p.1. Right: The relative reporter activity from the firefly luciferase in response to two miR-324 duplexes in HEK293T. The reporter activity from Renilla luciferase was used as a control. Bars indicate mean ± s.d. (n = 2, biological replicates). **p < 0.01 by the two-sided Student’s *t* test. **(C)** Cumulative distribution of changes in abundance of miR-324 targets after transfection of miR-324 duplexes in A172. Target genes were predicted by TargetScan (Agarwal et al., 2015). P-value by the one-sided Kolmogorov–Smirnov test. See also Figure S5.

This conforms well to the established strand selection rules. From the long duplex, the 5p strand is selected because the 5′ end of 5p is thermodynamically unstable compared to that of 3p.2 (Figure 5A, blue dashed rectangle). However, in the short duplex, 3p.1 starts with a 5′ adenosine which is favored by AGO (Figure 5A, red dashed rectangle). Taken together, the results demonstrate that alternative processing results in arm switching of miR-324.

### Alteration of the miR-324 strand ratio affects cell cycle progression

It was previously reported that the 5′ end of miR-324-3p varies in human glioblastoma (GBM) (Skalsky and Cullen, 2011). The proportion of 3p.1 is substantially greater in tumor than in normal brain tissue (Figure 6A). To examine the possibility that TUT4/7-mediated miR-324 maturation is differentially modulated in the GBM context, we analyzed publicly available transcriptome datasets. Both TUT4 and TUT7 are significantly upregulated in GBM (Figures 6B and S6A) (Lee and Choi, 2017; Madhavan et al., 2009). Using an independent dataset that profiled both mRNAs and miRNAs (Gulluoglu et al., 2018), we confirmed the upregulation of TUT4/7 levels in GBM and observed the reduction of the miR-324-5p/3p ratio in GBM compared to the matched normal samples (Figures 6C and S6B). It is noteworthy that TUT4/7 levels negatively correlate with 5p/3p ratios (Figure 6C). Moreover, GBM patients with low 5p/3p ratios exhibited poor prognosis (Figure S6C). These data collectively suggest that the TUT4/7-mediated miR-324 regulation may be physiologically relevant in GBM.

**Figure 6.**
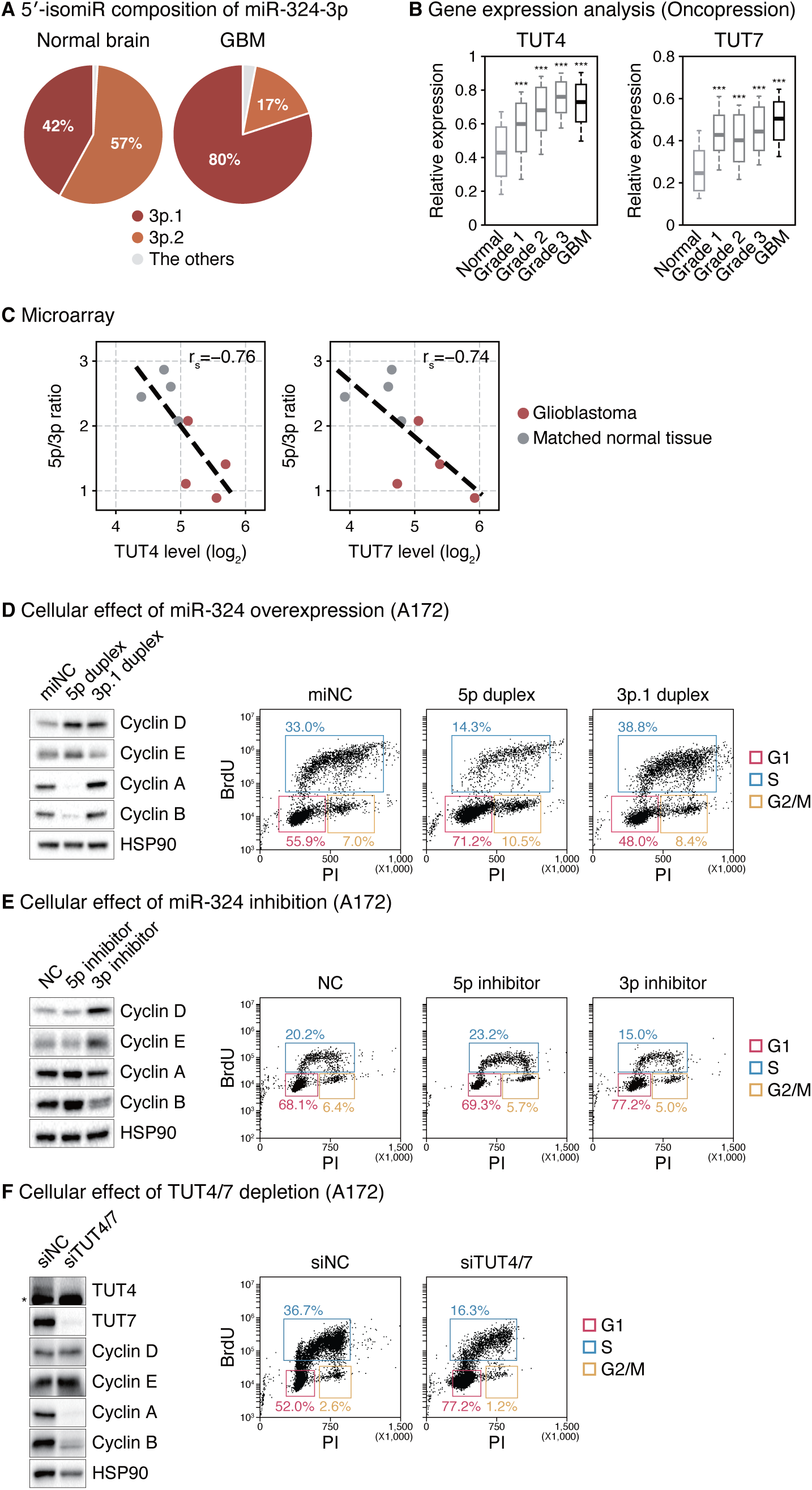
Alteration of the miR-324 strand ratio affects cell cycle progression. **(A)** 5′-isomiR composition of miR-324-3p in glioblastoma and normal brain tissues (Skalsky and Cullen, 2011). Normal brain, n = 3; glioblastoma, n = 6. **(B)** Expression levels of TUT4/7 in normal, lower grade glioma, and glioblastoma tissues from the Oncopression database. Normal, n = 723; grade 1, n = 74; grade 2, n = 133; grade 3, n = 132; glioblastoma, n = 865. ***p < 0.001 by one-way ANOVA with Tukey’s post hoc test for multiple comparisons. **(C)** Negative correlation between the TUT4/7 expression level and the miR-324-5p/3p ratio in glioblastoma (red) and matched normal brain tissues (grey) (Gulluoglu et al., 2018). The linear regression is shown with dashed lines. r_s_, Spearman correlation coefficient. **(D)** Western blot of indicated cyclin proteins and cell cycle profile after overexpressing synthetic 5p duplex (long duplex) or 3p.1 duplex (short duplex) of miR-324 in A172 cells. **(E)** Western blot of indicated cyclin proteins and cell cycle profile after inhibiting miR-324 by locked nucleic acid antisense oligonucleotides in A172 cells. **(F)** Western blot of indicated proteins and cell cycle profile after knocking down TUT4/7 in A172 cells. *, a cross-reacting band. PI, propidium iodide. BrdU, bromodeoxyuridine. See also Figure S6.

To understand the function of miR-324 arm switching, we investigated the molecular and cellular phenotypes driven by perturbation of the miR-324 strand selection in GBM cell lines. Given the previous studies showing functions of the miR-324 strands in cell proliferation in hepatocellular carcinoma (Cao et al., 2015; Tuo et al., 2017; Xiao et al., 2017), we examined the cell cycle. Transfection of the synthetic long duplex (“5p duplex”) or an antisense oligo against 3p (“3p inhibitor”) resulted in the dysregulation of several cyclin proteins; an accumulation of cyclins D and E and a reduction of cyclins A and B were observed (Figures 6D and 6E, left panels). Accordingly, cell cycle profiling indicated the cells are arrested at the G1 phase (Figures 6D and 6E, right panels). These molecular changes and cellular phenotypes were also observed with combinatorial treatment of the 5p duplex and the 3p inhibitor (Figures S6D and S6E). These observations are in line with previous studies where miR-324-5p reduces brain tumor growth in cultured cells and *in vivo* and that its downregulation in glioblastoma is associated with poor prognosis (Ferretti et al., 2008; Zhi et al., 2017). Lastly, TUT4/7 knockdown gave rise to cyclin dysregulation and G1 arrest in GBM cell lines (Figures 6F and S6F) and reduced cell viability in patient-derived GBM tumorsphere culture (Figure S6G). Together, these results indicate that miR-324 isomiRs have opposing functions in GBM cell proliferation.

## DISCUSSION

We learned from this study that approximately 14% of miRNAs examined undergo arm switching (Figure 1A). We further found that arm switching is an actively regulated process with defined *trans*-acting and *cis*-acting elements, at least in the case of miR-324 (Figure 7). In this pathway, TUT4/7 function as the key players by uridylating pre-miR-324. When modified, pre-miR-324 is cleaved at an alternative site by DICER, yielding a shorter duplex with a different end property. The shorter duplex is loaded onto AGO in an inverted orientation such that 3p.1 is selected instead of 5p. As the TUT4/7 levels are under a tissue/developmental stage-specific regulation, the arm ratio of miR-324 changes accordingly. TUT4/7 are also modulated under pathological conditions such as in glioblastoma, resulting in misregulation of miR-324.

**Figure 7.**
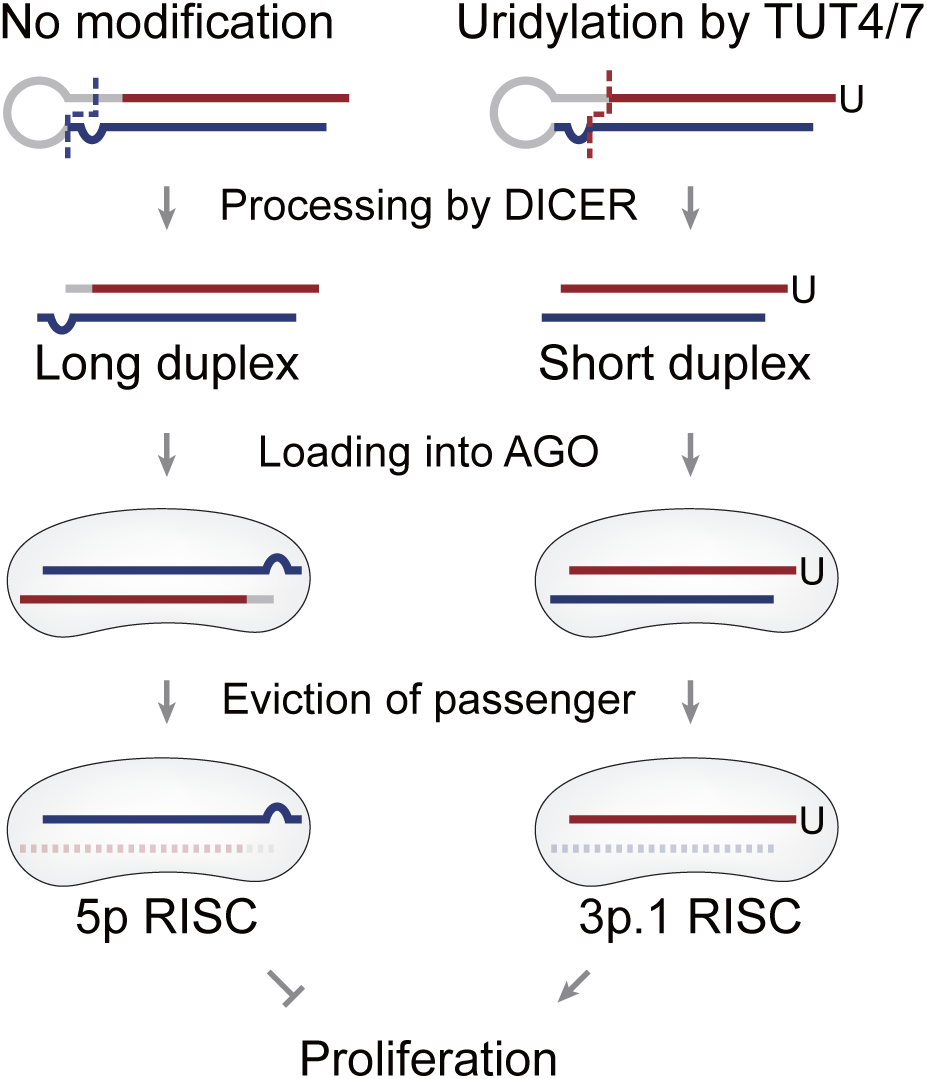
A proposed model of miR-324 arm switching mechanism and its functional consequences.

TUT4/7 uridylate many different RNA species including miRNAs, histone mRNAs, deadenylated mRNAs, viral RNAs, retrotransposon RNAs, rRNAs, vault RNAs, and Y RNAs (Hagan et al., 2009; Heo et al., 2012; Heo et al., 2009; Kim et al., 2015; Labno et al., 2016; Lackey et al., 2016; Le Pen et al., 2018; Lim et al., 2014; Pirouz et al., 2019; Thornton et al., 2012; Warkocki et al., 2018). Oligo-uridine tail recruits the 3′-5′ exoribonuclease DIS3L2 and thereby induces decay (Chang et Proliferation al., 2013). This decay mechanism is generally utilized to constantly remove undesired RNAs from the cytoplasm.

However, the miRNA pathway uses the uridylation machinery to achieve intricate and cell type-specific regulations. The most extensively studied example is the *let-7* biogenesis. TUT4/7 oligo-uridylate pre-*let-7* specifically in the presence of LIN28 that is expressed highly in early embryos and certain types of cancer (Hagan et al., 2009; Heo et al., 2008; Heo et al., 2009; Thornton et al., 2012). Oligo-uridylation blocks DICER processing and induces decay of pre-*let-7*. In the absence of LIN28, group II pre-*let-7* with the suboptimal 1-nt 3′ overhang structure is mono-uridylated by TUT4/7, which promotes DICER processing (Heo et al., 2012). Here, we reveal a new regulatory role for TUT4/7 in miRNA biogenesis. TUT4/7 mono- or di-uridylate pre-miR-324, which alters the DICER processing site and eventually leads to arm switching. Sequences and structure of pre-miR-324 are highly conserved in mammals (Figure S1D). Furthermore, the alternative isoforms are observed in all mammalian datasets (Figure 1D), supporting the biological relevance of TUT4/7-mediated alternative maturation of this miRNA.

*De novo* identification of isomiRs using sRNA-seq helped us to gain molecular insights of arm switching (Figure 1B). Comparative sRNA-seq analyses revealed many miRNAs with varying isomiR levels, which warrants future investigation. Figure S7 presents some interesting cases whose strand ratio and 5′-isomiR fraction fluctuate depending on cell types. For instance, miR-296, miR-671, and miR-676 show highly variable isomiR profiles. Thus, we anticipate that regulated arm switching is not limited to miR-324. Of note, we have not found evidence for uridylation on these miRNAs, suggesting different mechanism(s) may be at work. Alternative cleavage by DROSHA or DICER may partially explain the arm switching events because it can produce miRNA duplexes with different termini (Chiang et al., 2010; Kim et al., 2017; Kim et al., 2019; Ma et al., 2016; Tan et al., 2014; Wu et al., 2009). In addition, a recent study reported that miRNA arm ratio can be influenced in part by differential decay rates which involves target-directed miRNA degradation (TDMD) (Zhang et al., 2019). It is unclear if and to what extent alternative processing and TDMD explain the variations we observe with sRNA-seq analyses across cell types/development (Figure S7). Future studies will be needed to elucidate the molecular mechanisms and physiological significance of regulation of these miRNAs.

In this study, we delineate the molecular mechanism of alternative DICER processing and revealed the role of the dsRBD. The DICER dsRBD has been known as an auxiliary domain that increases the overall RNA binding affinity of DICER and enhances processing efficiency (Ma et al., 2012; Zhang et al., 2004). However, our new results demonstrate that the dsRBD repositions pre-miR-324 in DICER, leading to a shift in the processing site. Thus, the dsRBD determines the processing sites together with the platform and PAZ domains that harbor the 5′ and 3′ pockets. This is interesting in light of our recent finding that the dsRBD of DROSHA also contributes to cleavage site selection by recognizing the “mGHG motif” on pri-miRNA (Fang and Bartel, 2015; Kwon et al., 2019). This role of the dsRBD may be deeply conserved across RNase III family members as the class I RNase III enzymes from *Aquifex aeolicus* and *Escherichia coli* also use the dsRBD for specific recognition of a sequence motif (Blaszczyk et al., 2004; Gan et al., 2006; Pertzev and Nicholson, 2006; Shi et al., 2011). It would be interesting to investigate to what extent the dsRBD is involved in the fidelity of DICER processing and which consensus sequence is recognized by the DICER dsRBD. Previously reported structures of human RNase III DROSHA and DICER do not contain the substrate RNAs in the processing-competent conformation (Kwon et al., 2016; Liu et al., 2018). Further structural and biochemical studies will be needed to understand the basis of this deeply conserved yet class-specific substrate recognition mechanism.

Lastly, we demonstrate that in GBM, the upregulated TUT4/7 control the miR-324 maturation in favor of 3p.1. Intriguingly, similar regulation may be in effect in hepatocellular carcinoma (HCC) where miR-324-5p is reduced and miR-324-3p.1 is upregulated (Cao et al., 2015; Tuo et al., 2017; Xiao et al., 2017). Previous studies collectively suggest that miR-324-5p and miR-324-3p.1 have divergent roles in HCC progression. It remains to be investigated if miR-324 arm switching in HCC is also mediated by TUT4/7. It would be useful to examine what other types of diseases are associated with dysregulation of TUT4/7 and miRNA arm switching (Kuo et al., 2016). Developing chemical inhibitors against TUT4/7 and testing their effects on tumor would also be of interest. Moreover, because the ratios between miR-324 isoforms differ depending on cell conditions/types, the miR-324 arm ratio may serve as a sensitive reporter for TUT4/7 activity *in vivo* and a marker for cancer diagnosis. Our study provides insights into the biological roles and potential applications of miRNA regulation and modification.

## METHODS

### AQ-seq library preparation

AQ-seq (bias-minimized sRNA-seq) libraries were constructed using total RNAs from fifteen mouse tissues (Mouse Total RNA Master Panel; Takara) as described in our previous study (Kim et al., 2019). Briefly, we mixed 5 µg of total RNAs with 10 fmole of thirty equimolar spike-in RNAs which are miRNA-like non-human/mouse/frog/fish RNAs used for bias evaluation. Small RNAs were enriched by size fractionation by 15% urea-polyacrylamide gel electrophoresis and sequentially ligated to the randomized adapter at the 3′ and 5′ ends. The ligated RNAs were reverse-transcribed using SuperScript III reverse transcriptase (Invitrogen), amplified using Phusion High-Fidelity DNA Polymerase (Thermo Scientific), and subjected to high-throughput sequencing on the MiSeq platform (Illumina).

### Analysis of high-throughput sequencing of miRNAs

Data processing was performed as described in our previous study (Kim et al., 2019) except that reads of the sRNA-seq results from mouse were mapped to the mouse genome (mm10). Briefly, the 3′ adapter was clipped from the reads using cutadapt (Martin, 2011). For AQ-seq data, 4-nt degenerate sequences were further removed with FASTX-Toolkit (http://hannonlab.cshl.edu/fastx_toolkit/). After filtering out short, low-quality, and artifact reads using FASTX-Toolkit, AQ-seq data were aligned to the spike-in sequences first and the unaligned reads were mapped to the genome next, while the other sRNA-seq data were mapped to the genome using BWA (Li and Durbin, 2010). For a given read, we selected alignment result(s) with the best alignment score allowing mismatches only at the 3′ end. miRNA annotations were retrieved using miRBase release 21 by the intersect tool in BEDTools (Kozomara and Griffiths-Jones, 2014; Quinlan and Hall, 2010).

For quantitative analysis of miRNA strand ratios, we first identified the most abundant 5′-isomiR for a given mature miRNA in the most abundantly expressed cell line or tissue. Then, the ratio between the top 5′-isomiRs from 5p and 3p was calculated for all of the cell lines or tissues. Non-repetitive miRNA genes whose both strands are annotated in miRBase release 21 were included in this analysis.

### Targetome analysis

Results from two modified versions of AGO CLIP-seq (CLEAR-CLIP and CLASH) were analyzed to identify miR-324 targets (Helwak et al., 2013; Moore et al., 2015). For analysis of CLEAR-CLIP, the 3′ and 5′ adapters were clipped using cutadapt followed by extraction of reads containing miR-324 sequences. The target RNA sequences were then mapped to the mouse genome (mm10). Annotations were retrieved using GENCODE release M19 by the intersect tool in BEDTools. For analysis of CLASH, the supplementary file of the CLASH data generated by the protocol E4 was used. Target genes of miR-324-3p were subdivided according to the 5′ end of the miR-324-3p sequences.

For prediction of miR-324 targets, we utilized TargetScan Human release 7.2 where target sites were identified based on the seed sequences of miR-324-5p (GCAUCC) or miR-324-3p.1 (CUGCCC) (Agarwal et al., 2015). Predicted targets with cumulative weighted context++ score below −0.2 were included in this analysis.

### Plasmid construction

To construct plasmids for expression of wild type, 5′ pocket mutant, and 3′ pocket mutant DICER, the coding sequence of the current human textitDICER reference (RefSeq NM_030621) was amplified with or without mutations introduced in our previous study (Park et al., 2011). The amplified DNAs were subcloned into the pCK-FLAG vector (CMV promoter-driven vector) at the BamHI and XhoI sites using In-Fusion HD Cloning Kit (Clontech). For dsRBD-deleted DICER, the coding region except the sequence corresponding to V1849–S1922 amino acids was amplified and subcloned into the pCK-FLAG vector in the same way. For TRBP, the coding sequence of the current human textitTRBP reference (RefSeq NM_134323) was amplified and subcloned into the pcDNA3-cMyc vector (Invitrogen) at the BamHI and XhoI sites. To construct plasmids for luciferase assay, synthetic DNA oligos containing three 8mer target sites of either miR-1-3p, miR-324-5p, or miR-324-3p.1 were amplified and inserted into pmirGLO (Promega) at the XhoI and XbaI sites. The sequences of synthetic DNA oligos and PCR primers are listed in Supplementary Table S1.

### Cell culture and transfection

A172 and U87MG were obtained from Korean Cell Line Bank. HEK293T, A172, and U87MG were maintained in DMEM (WELGENE) supplemented with 10% fetal bovine serum (WELGENE). Primary tumor cells derived from a glioblastoma patient (TS13-64) were established from fresh glioblastoma tissue specimens, as approved by the institutional review board of Yonsei University College of Medicine (4-2012-0212, 4-2014-0649). For tumorsphere culture, TS13-64 cells were grown in DMEM (WELGENE) supplemented with 1X B-27 (Thermo Scientific), 20 ng/mL of bFGF (R&D Systems), and 20 ng/mL of EGF (Sigma-Aldrich) (Kong et al., 2013).

To knock down TUT4/7, cells were transfected with 20–22 nM siRNAs using the Lipofectamine 3000 reagent (Thermo Scientific) twice and harvested 4 days after 1st transfection. For overexpression of DICER, HEK293T cells were transfected with pCK-FLAG-DICER using the Lipofectamine 3000 reagent (Thermo Scientific) and harvested 2 days post transfection. To deliver synthetic miRNAs or inhibitors, cells were transfected with 20 nM of synthetic miRNA duplexes or 40 nM of LNA miRNA inhibitors using Lipofectamine 3000 reagent (Thermo Scientific) and harvested 2 days after transfection. For simultaneous transfection of synthetic miRNAs and inhibitors, cells were transfected with 20 nM of synthetic miRNA duplexes and 80 nM of LNA miRNA inhibitors. For RNA-seq, we harvested cells 1 day after transfection to minimize the secondary effect.

Control siRNA (AccuTarget Negative Control siRNA), siRNAs, control miRNA, and synthetic miR-324 duplexes were obtained from Bioneer. Control miRNA inhibitor (miRCURY LNA miRNA Inhibitor Negative control A) and miR-324 inhibitors (hsa-miR-324-5p and hsa-miR-324-3p miRCURY LNA miRNA Inhibitor) were obtained from QIAGEN. The sequences of synthetic siRNAs and miRNAs are listed in Supplementary Table S1.

### Northern blot

Total RNAs were isolated using TRIzol (Invitrogen) and small RNAs (< 200 nt) were enriched with the mirVana miRNA Isolation Kit (Ambion). The RNAs were resolved on 15% urea-polyacrylamide gels. Synthetic miR-324 duplex and Decade Markers System (Ambion) were loaded as size markers. RNAs were transferred to Hybond-NX membranes (Amersham) and crosslinked to the membranes with 1-ethyl-3-(3-dimethylaminopropyl) carbodiimide (Pall and Hamilton, 2008). Antisense probes were radiolabeled at their 5′ ends with [*γ*-^32^P] ATP by T4 polynucleotide (Takara) and purified using Performa Spin Columns (Edge BioSystems). The membranes were incubated with denatured UltraPure Salmon Sperm DNA Solution (Thermo Scientific) in PerfectHyb Plus Hybridization Buffer, hybridized with antisense probes, and washed with mild wash buffer (0.05% SDS and 2X SSC) followed by stringent wash buffer (0.1% SDS and 0.1X SSC). Radioactive signals were detected by Typhoon FLA 7000 (GE Healthcare) and analyzed using the Multi Gauge software (FujiFilm). miR-324-3p was detected first and miR-324-5p was detected next after stripping off the probes. Lastly, U6 snRNA was detected after stripping off the probes. To remove the probes from the blot, the membrane was soaked in pre-boiled 0.5% SDS for 15 min. Synthetic miR-324 duplex (AccuTarget) was obtained from Bioneer. The sequences of probes are listed in Supplementary Table S1.

### Quantitative real-time PCR (RT-qPCR)

To measure mRNA levels in mouse tissues, RNAs from Mouse Total RNA Master Panel (Takara) were reverse-transcribed using the RevertAid First Strand cDNA Synthesis Kit (Thermo Scientific) and subjected to quantitative real-time PCR with the Power SYBR Green PCR Master Mix (Thermo Scientific) on StepOnePlus Real-Time PCR System (Thermo Scientific). GAPDH was used for internal control. The sequences of qPCR primers are listed in Supplementary Table S1.

To quantify the miR-324-5p/3p strand ratio, total RNAs were isolated using TRIzol (Invitrogen). cDNAs were then synthesized using the TaqMan miRNA Reverse Transcription kit (Applied Biosystems) and subjected to quantitative real-time PCR with the TaqMan gene expression assay kit (Applied Biosystems) on StepOnePlus Real-Time PCR System (Thermo Scientific). U6 snRNA was used for internal control.

### *In vitro* DICER processing assay

Pre-miR-324 and its variants were prepared by ligating two synthetic RNA fragments as previously described (Heo et al., 2009). They were radiolabeled at their 5′ ends with [*γ*-^32^P] ATP by T4 polynucleotide (Takara) and purified using Oligo Clean & Concentrator (Zymo Research) according to manufacturer’s instructions.

For immunoprecipitation of FLAG-DICER, the HEK293T cells overexpressing DICER proteins were lysed with lysis buffer (500 mM NaCl, 20 mM Tris (pH 8.0), 1 mM EDTA, 1% Triton X-100) and subjected to sonication using Bioruptor Standard (Diagenode). After centrifugation, the supernatant was incubated with 10 µL of ANTI-FLAG M2 Affinity Gel (Sigma-Aldrich). The beads were washed twice with lysis buffer, four times with high salt buffer (800 mM NaCl and 50 mM Tris (pH 8.0)), and four times with buffer D (200 mM KCl, 20 mM Tris (pH 8.0), 0.2 mM EDTA) and then resuspended in 10 µL buffer D.

The immunopurified DICER was subjected to *in vitro* reactions in a total volume of 30 µL containing 2 mM MgCl_2_, 1 mM DTT, 100 mM KCl, 10 mM Tris (pH 8.0), 0.1 mM EDTA, 60 units of SUPERase In RNase Inhibitor (Thermo Scientific), and the 5′-radiolabeled pre-miR-324. The RNAs were purified with phenol extraction or Oligo Clean & Concentrator (Zymo Research) and resolved on 15% urea-polyacrylamide gels. Synthetic miR-324-5p and Decade Markers System (Ambion) were loaded as size markers. Synthetic pre-miR-324 fragments were obtained from IDT. Synthetic miR-324-5p was obtained from Bioneer. The sequences of synthetic pre-miR-324 fragments and miR-324-5p are listed in Supplementary Table S1.

### Dual luciferase reporter assay

HEK293T cells were co-transfected with pmirGLO containing three 8mer target sites of either miR-1-3p, miR-324-5p, or miR-324-3p.1 along with control miRNA or miR-324 duplexes using the Lipofectamine 3000 reagent (Thermo Scientific). After 2 days of transfection, cells were harvested and subjected to reporter assay. The reporter activities were measured using Dual-Luciferase Reporter Assay System according to the manufacturer’s instructions (Promega) on the Spark microplate reader (TECAN). Given that miR-1-3p is little expressed in HEK293T (∼8 RPM in the AQ-seq result) (Kim et al., 2019), pmirGLO with three 8mer target sites of miR-1-3p was used for a plasmid control. Control miRNA was used for further normalization.

### RNA-seq

A172 cells were transfected with control miRNA or miR-324 duplexes using the Lipofectamine 3000 reagent (Thermo Scientific). Cells were harvested after 1 day of transfection and total RNAs were isolated using TRIzol (Invitrogen). The RNA quality was confirmed using Agilent 2100 Bioanalyzer and DNA nanoball sequencing for transcriptome (oligo dT enrichment; stranded; 100 bp paired-end) was performed on the BGISEQ-500 platform by BGI Tech Solutions (Hong Kong).

For gene expression analysis, references were first built with the primary assembly from the human genome and “BestRefSeq” and “Curated Genomic” sources from the annotation of the RefSeq assembly accession GCF_000001405.34 using RSEM with “–bowtie2” options (Li and Dewey, 2011). Then, RNA-seq data were analyzed to estimate gene expression levels using RSEM with “–paired-end” and “–bowtie2” options.

### Gene expression analysis of glioblastoma patients data

From Oncopression (http://oncopression.com), preprocessed gene expression data using microarray were retrieved (Lee and Choi, 2017). The REMBRANDT gene expression dataset (E-MTAB-3073) was obtained from ArrayExpress (Madhavan et al., 2009). For analysis of simultaneous profiling of mRNAs and miRNAs (Gulluoglu et al., 2018), raw microarray data were normalized by the robust multi-array average (RMA) using the limma package in R and then used for gene expression analysis.

### Survival analysis

The level 3 miRNA gene quantification data and clinical data for TCGA glioblastoma patients were obtained from the GDC legacy archive and the GDC data portal, respectively. The patients were stratified according to the miR-324-5p/3p ratio and the top and bottom 40% of the cases were included in the analysis. The patient’s survival was estimated by the Kaplan-Meier method and tested by the two-sided log-rank test using the survival package in R.

### Western blot

Cells were harvested, washed with PBS and lysed in RIPA buffer (Thermo Scientific) complemented with protease inhibitor cocktail set III (Merck Millipore) and phosphatase inhibitor cocktail II (AG Scientific). Protein concentration was measured with the BCA Protein Assay Kit (Pierce Biotechnology) and equal amounts of protein were separated on 4–12% Tris-Glycine Gels (Thermo Scientific) and transferred onto Immobilon-P PVDF membranes (Merck Millipore). Membranes were incubated with 5% skimmed milk in PBS-T (PBS (Amresco) + 0.1% Tween 20 (Anatrace)) and then probed with primary antibodies. After being washed three times with PBS-T, the membranes were incubated with HRP-conjugated secondary antibodies. The protein bands were detected by SuperSignal West Pico PLUS Chemiluminescent Substrate (Thermo Scientific) and scanned by ChemiDoc XRS+ System (Bio-Rad).

#### Antibodies

The rabbit polyclonal antibodies against TUT4 (18980-1-AP, RRID:AB_10598327, 1:500) and TUT7 (25196-1-AP, 1:500) were purchased from Proteintech. The mouse monoclonal antibody against Cyclin E (sc-247, RRID:AB_627357, 1:1,000) and rabbit polyclonal antibodies against Cyclin A (sc-751, RRID:AB_631329, 1:1,000), Cyclin B1 (sc-752, RRID:AB_2072134, 1:1,000), and Cyclin D1 (sc-753, RRID:AB_2070433, 1:1,000) were purchased from Santa Cruz Biotechnology. The rabbit polyclonal antibody against HSP90 (4874, RRID:AB_2121214, 1:1,000) was purchased from Cell Signaling. The HRP-conjugated goat polyclonal antibodies against rabbit IgG (111-035-144, RRID:AB_2307391, 1:10,000) and mouse IgG (115-035-146, RRID:AB_2307392, 1:10,000) were purchased from Jackson ImmunoResearch.

### Flow cytometry

Cells were incubated with 10 µM BrdU for 3–8 hr before fixation by ice-cold 70% ethanol. The fixed cells were incubated with FITC-conjugated anti-BrdU antibody (11-5071-42, RRID:AB_11042627, Invitrogen), further stained with 20 µg/mL of propidium iodide (Sigma-Aldrich) in the presence of 10 µg/mL of RNase A (Thermo Scientific), and detected using BD Accuri C6 Plus Flow Cytometer. Cell cycle was analyzed by the BD Accuri C6 system software.

## SUPPLEMENTAL INFORMATION

**Supplementary Table 1**

Lists of oligonucleotide sequences used in this study.

## ACKNOWLEDGMENTS

We are grateful to Eunji Kim for cloning and mutagenesis of expression constructs. We also thank Yoonseok Jung, Dongwan Kim, Hyunjoon Kim, Kijun Kim, Myeonghwan Kim, Sung-Chul Kwon, Yongwoo Na, Jae Ho Paek, Soomin Son, Buyeon Um, Hyerim Yi, and other members of our laboratory for their helpful discussions. The survival analysis result is based upon data generated by the TCGA Research Network: https://www.cancer.gov/tcga. This research was supported by the Institute for Basic Science from the Ministry of Science and ICT of Korea (IBS-R008-D1 to H.K., J.K., S.Y., Y.Y.L., and V.N.K.), the BK21 Research Fellowships from the Ministry of Education of Korea (to H.K. and Y.Y.L.), and the NRF (National Research Foundation of Korea) Grant funded by the Korean government (NRF-2015-Global Ph.D. Fellowship Program to H.K. and NRF-2018-Global Ph.D. Fellowship Program to Y.-Y.L.).

## AUTHOR CONTRIBUTIONS

Conceptualization, H.K., J.K., and V.N.K.; Methodology, H.K., J.K., S.Y., Y.-Y.L., and V.N.K.; Formal Analysis, H.K., J.P., R.J.C., and S.J.Y.; Investigation, H.K., J.K., S.Y., Y.-Y.L., J.P., R.J.C., and S.J.Y.; Resources, J.P., R.J.C., S.J.Y., and S.-G.K.; Writing – Original Draft, H.K., J.K., S.Y., J.P., R.J.C., S.J.Y., S.-G.K., and V.N.K.; Writing –Review & Editing, H.K., J.K., S.Y., Y.-Y.L., and V.N.K.; Visualization, H.K., J.K., and V.N.K.; Supervision, S.-G.K. and V.N.K.; Funding Acquisition, H.K., Y.-Y.L., and V.N.K.;

## DECLARATION OF INTERESTS

The authors declare no competing interests.

---

**Figure S1, related to Figure 1.**
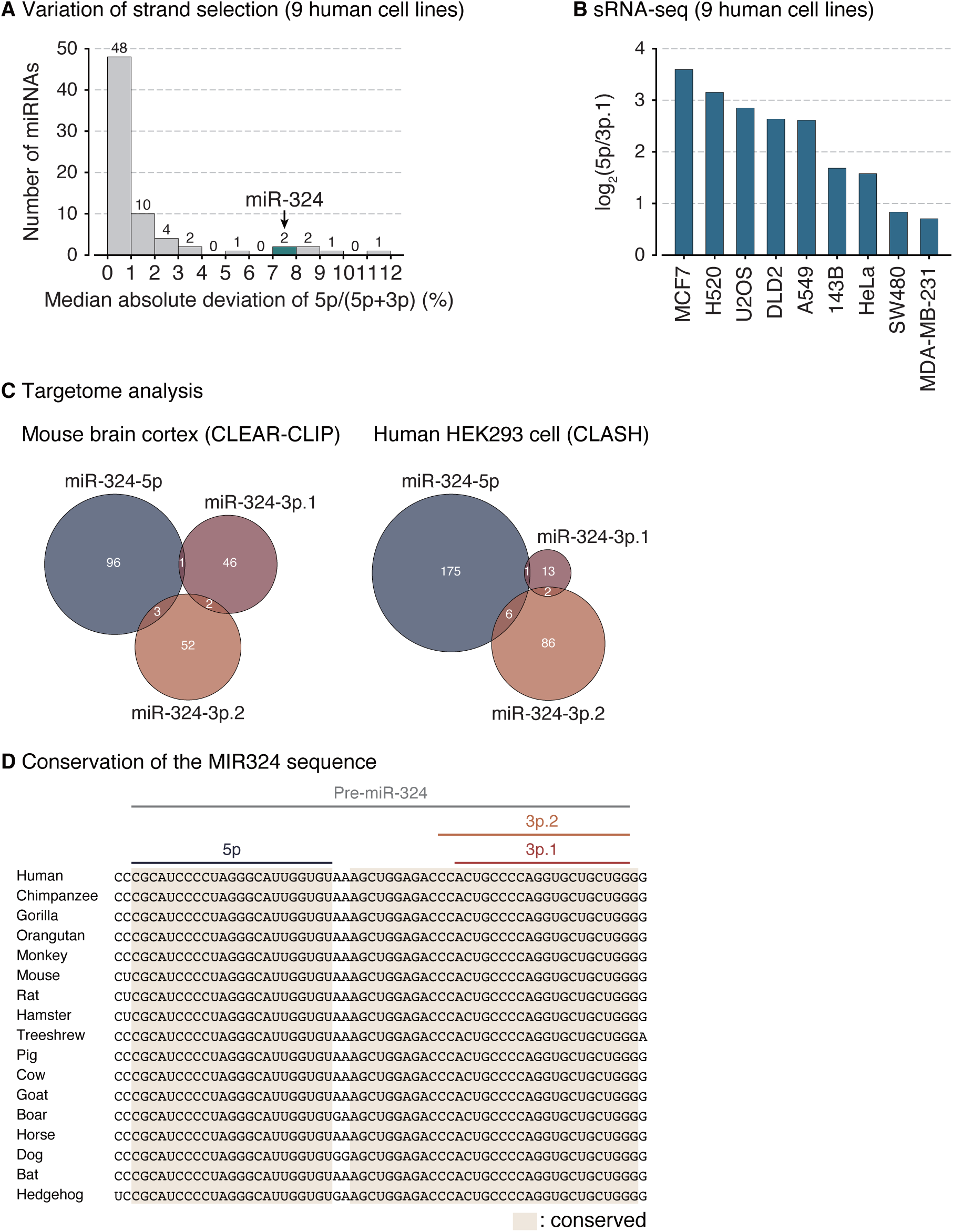
miR-324 undergoes alternative strand selection and has conserved and functionally distinct 5′-isomiRs. **(A)** Median absolute deviation of the 5p proportion for a given miRNA in human cell lines. Abundant and ubiquitous miRNAs (> 100 median RPM in both human cell line and mouse tissue datasets, > 0 RPM in all samples) were included in this analysis. **(B)** Strand ratios (5p/3p.1) of miR-324 across the indicated panel of human cell lines as measured by sRNA-seq. **(C)** Targetome analysis of three 5′-isomiRs of miR-324. Target RNAs were identified from chimeric reads composed of each 5′-isomiR and bound target RNAs. **(D)** Sequence conservation of the MIR324 gene in mammals.

**Figure S2, related to Figure 2.**
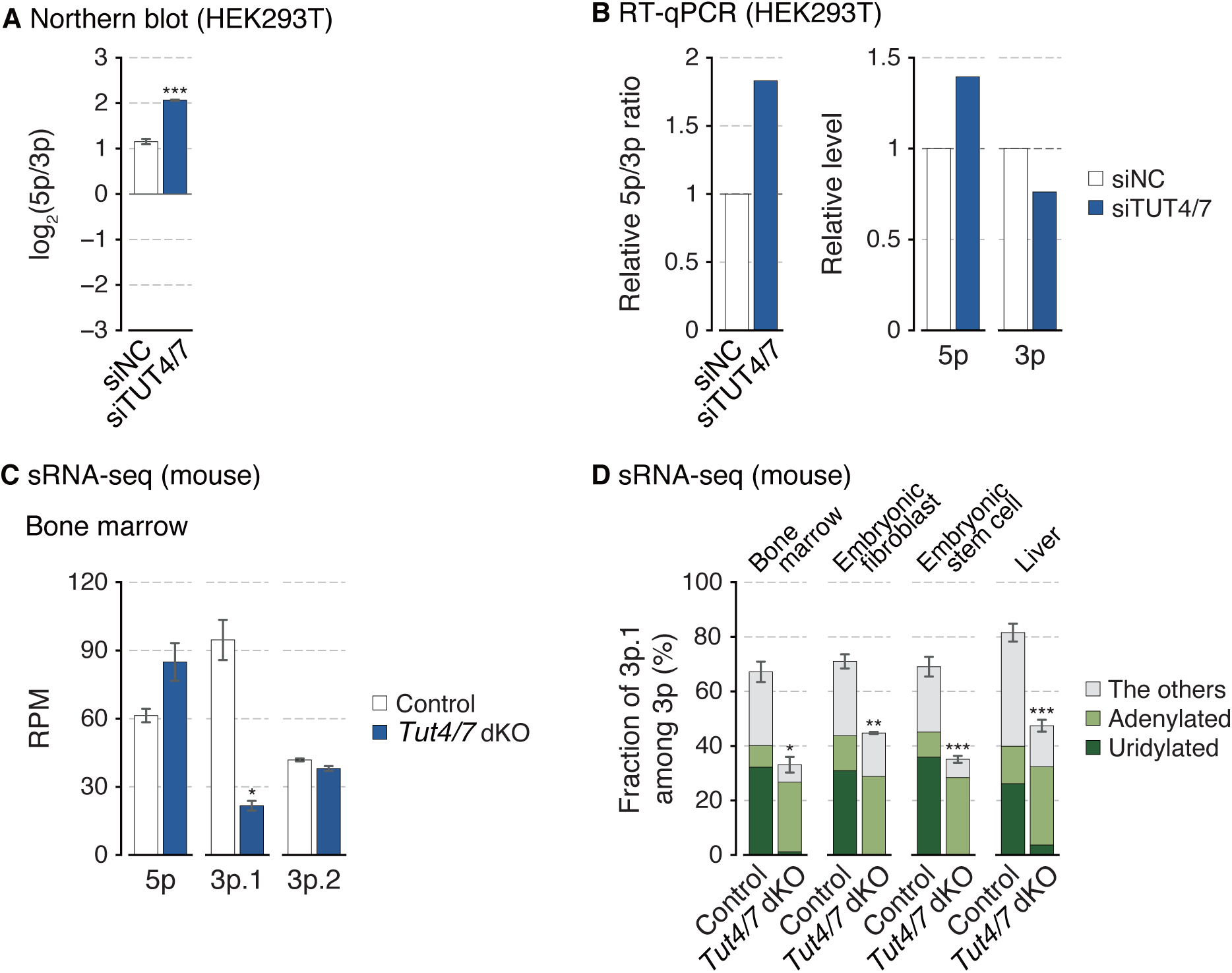
TUT4/7 regulate miR-324 arm switching. **(A)** The miR-324-5p/3p ratio calculated from band intensities of the northern blot (Figure 2D). Synthetic miR-324 duplex was used for normalization. Bars indicate mean ± s.d (n = 3, biological replicates). ***p < 0.001 by the two-sided paired *t* test. **(B)** The relative miR-324-5p/3p ratio and their expression levels measured by RT-qPCR after knockdown of TUT4/7 in HEK293T. **(C)** Relative abundance of three 5′-isomiRs of miR-324 in mouse bone marrow. Bars indicate mean ± s.d. (n = 2, biological replicates). *p < 0.05 by the two-sided Student’s *t* test. RPM, reads per million. **(D)** Fraction of miR-324-3p.1 among 3p strand in mice. The type of the 3′ end modification is indicated with different colors. Bars indicate mean ± s.d. (bone marrow, n = 2; embryonic fibroblast, n = 2; embryonic stem cell, n = 3; control and *Tut4/7* dKO in liver, n = 4 and 3, respectively). *p < 0.05, **p < 0.01, ***p < 0.001 by the two-sided Student’s *t* test.

**Figure S3, related to Figure 3.**
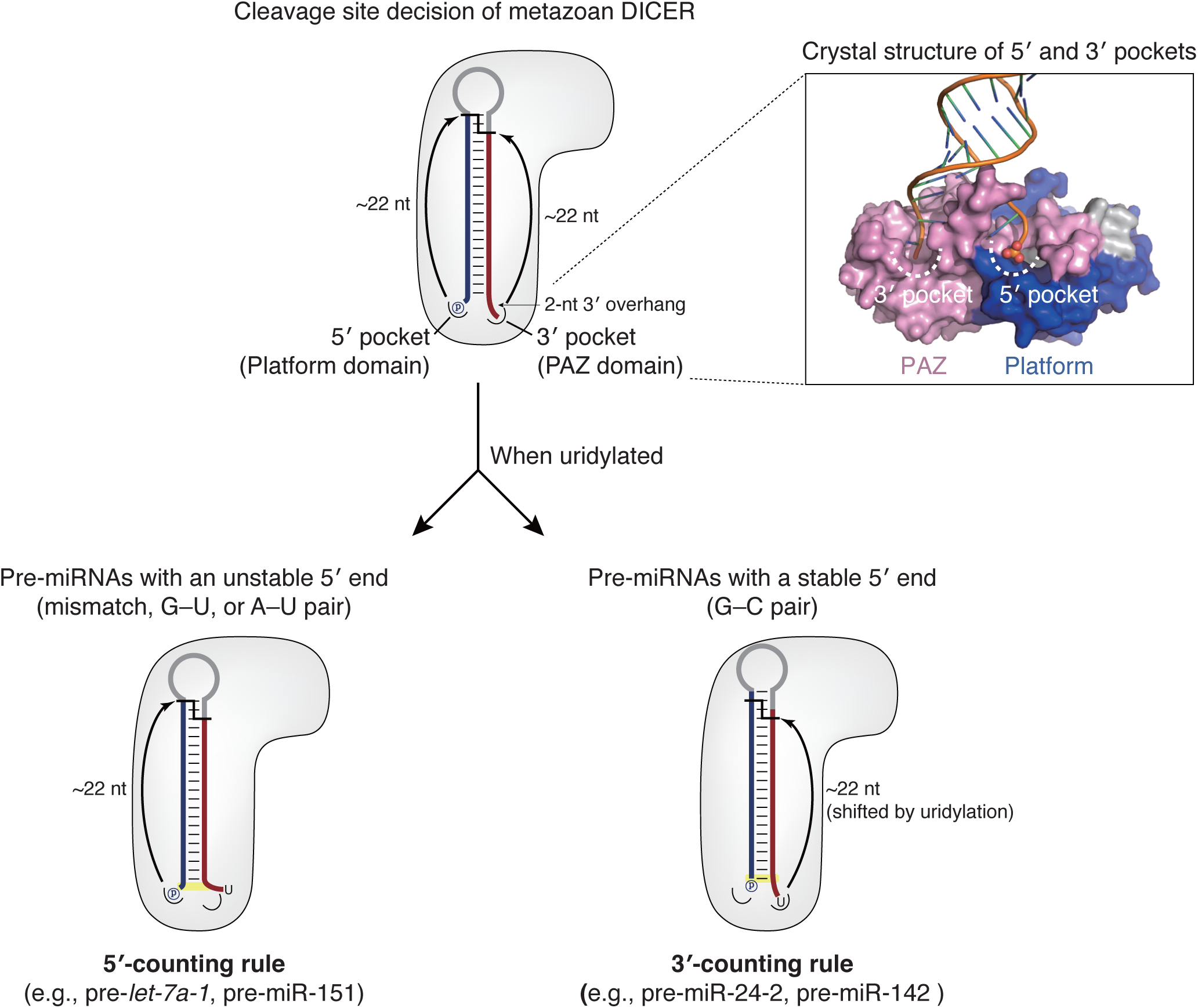
DICER has been known to decide the cleavage site by measuring ∼22 nt from the 5′ or 3′ ends. In the crystal structure, the platform domain and the PAZ domain are colored in blue and pink, respectively (Protein Data Bank (PDB) ID 4NH5) (Tian et al., 2014).

**Figure S4, related to Figure 4.**
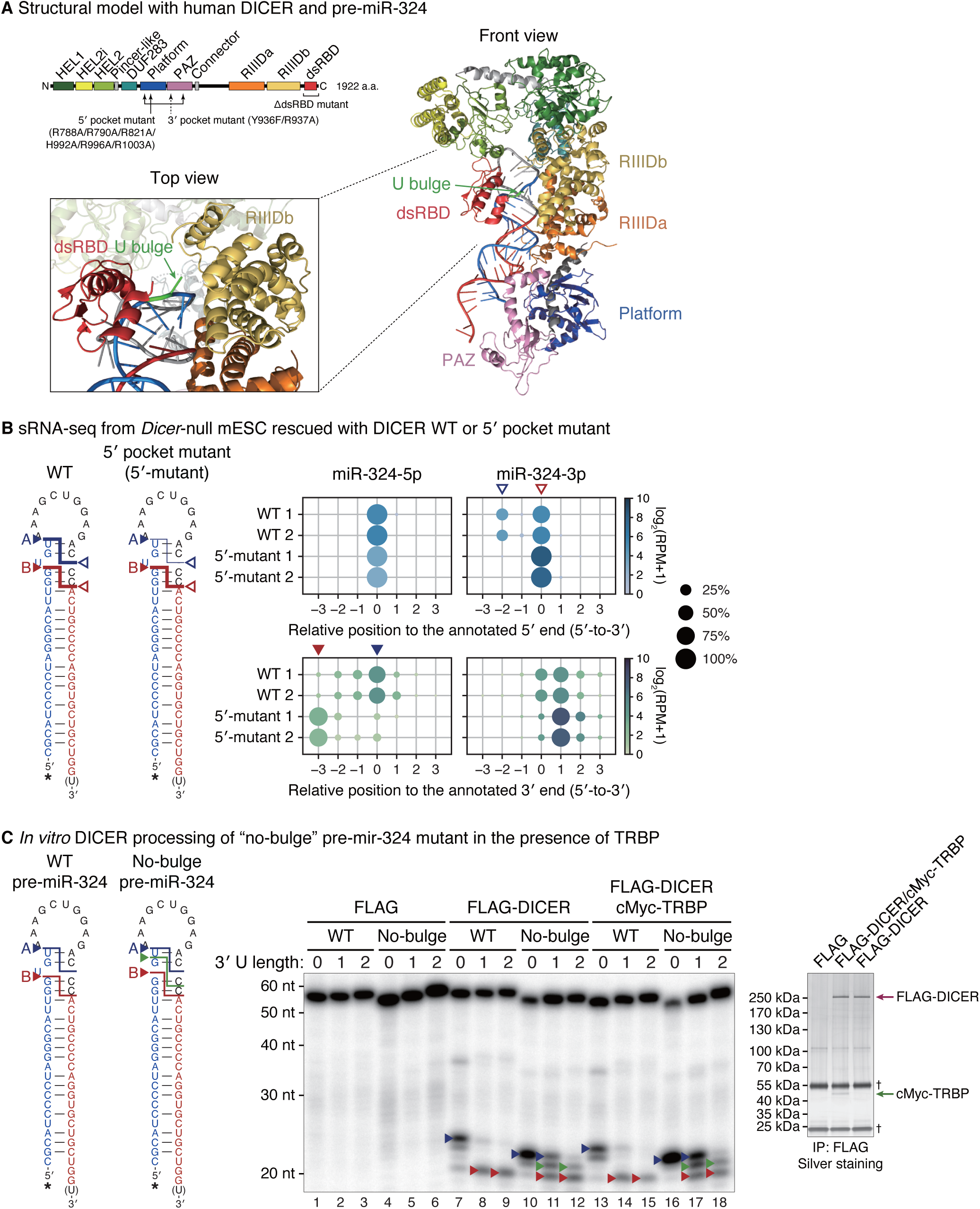
Alternative DICER processing site is not dependent on the 5′-counting rule but dictated by the U bulge structure of pre-miR-324. **(A)** The structural model of human DICER and pre-miR-324. The orientation of unmodified pre-miR-324 on human DICER was modeled, based on the crystal structure of double-stranded RNA-bound *Aquifex aeolicus* (Aa) RNase III (Gan et al., 2006; Liu et al., 2018). The DICER dsRBD was then superimposed with that of Aa RNase III to predict its position in a dicing state. The U bulge of pre-miR-324 is in the vicinity of RIIIDb in the model. HEL1, HEL2i, and HEL2 in the DExD/H-box helicase domain in green, lemon, and yellow-green, respectively; the DUF283 domain in cyan; the platform domain in blue, the PAZ domain in pink; RIIIDa and RIIIDb, the RNase IIIa and RNase IIIb domains in orange and yellow, respectively; dsRBD, the double-stranded RNA binding domain in red. **(B)** The usage of the indicated 5′ and 3′ ends of miR-324 in the *Dicer*-null mouse embryonic stem cells (mESCs) rescued with wild type or 5′ pocket mutant DICER. The 5′ and 3′ ends of human miR-324 annotated in miRBase release 21 were used as references. Cleavage sites and their corresponding positions on miR-324 are marked with arrowheads. RPM, reads per million. **(C)** Left: *In vitro* processing of wild type or no-bulge mutant pre-miR-324 by immunopurified DICER along with co-immunoprecipitated TRBP. Lanes 1–12 (FLAG and FLAG-DICER) are identical with those in Figure 4A. Major cleavage products and their corresponding cleavage sites are marked with arrowheads. *, radiolabeled 5′ phosphates. Right: Immunopurified proteins analyzed by silver staining. †, heavy and light chains of anti-FLAG antibodies.

**Figure S5, related to Figure 5.**
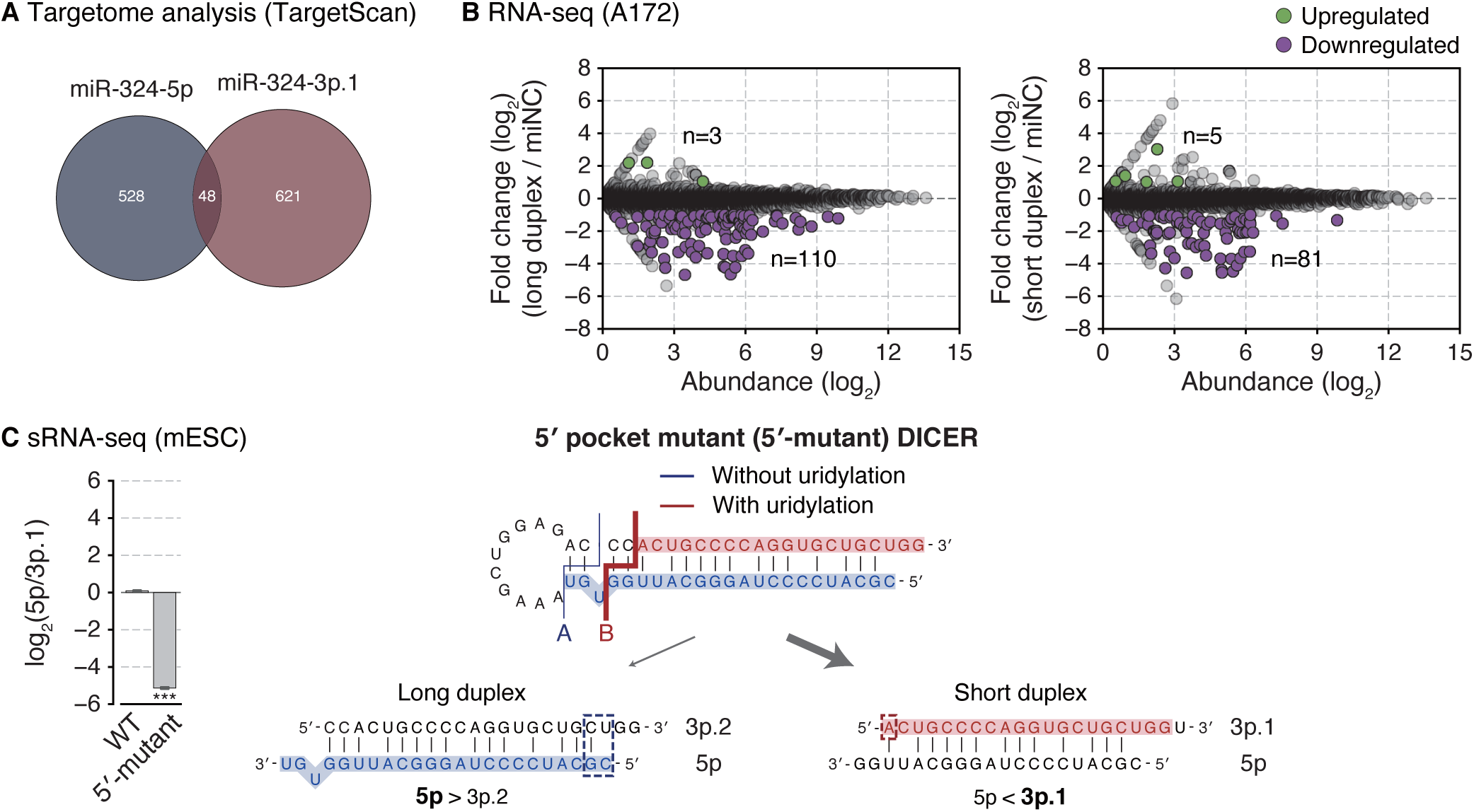
Alternative DICER processing is responsible for the arm switching. **(A)** Targetome analysis of miR-324-5p and 3p. Among the genes expressed in A172, targets were predicted by TargetScan. **(B)** Gene expression changes upon transfection of miR-324 duplexes in A172. Differentially expressed genes (p-value < 0.001 by edgeR (Robinson et al., 2010)) with log_2_-transformed fold change > 1 or < −1 were indicated with green or purple dots, respectively. **(C)** Strand selection of miR-324 in the *Dicer*-null mESCs replenished with wild type or 5′ pocket mutant DICER. Left: The miR-324-5p/3p.1 ratio detected by sRNA-seq. Bars indicate mean ± s.d. (n = 2, biological replicates). ***p < 0.001 by the two-sided Student’s *t* test. Right: A schematic diagram of strand selection of miR-324 duplexes generated from the processing by 5′ pocket mutant DICER (Figures 4B and S4B).

**Figure S6, related to Figure 6.**
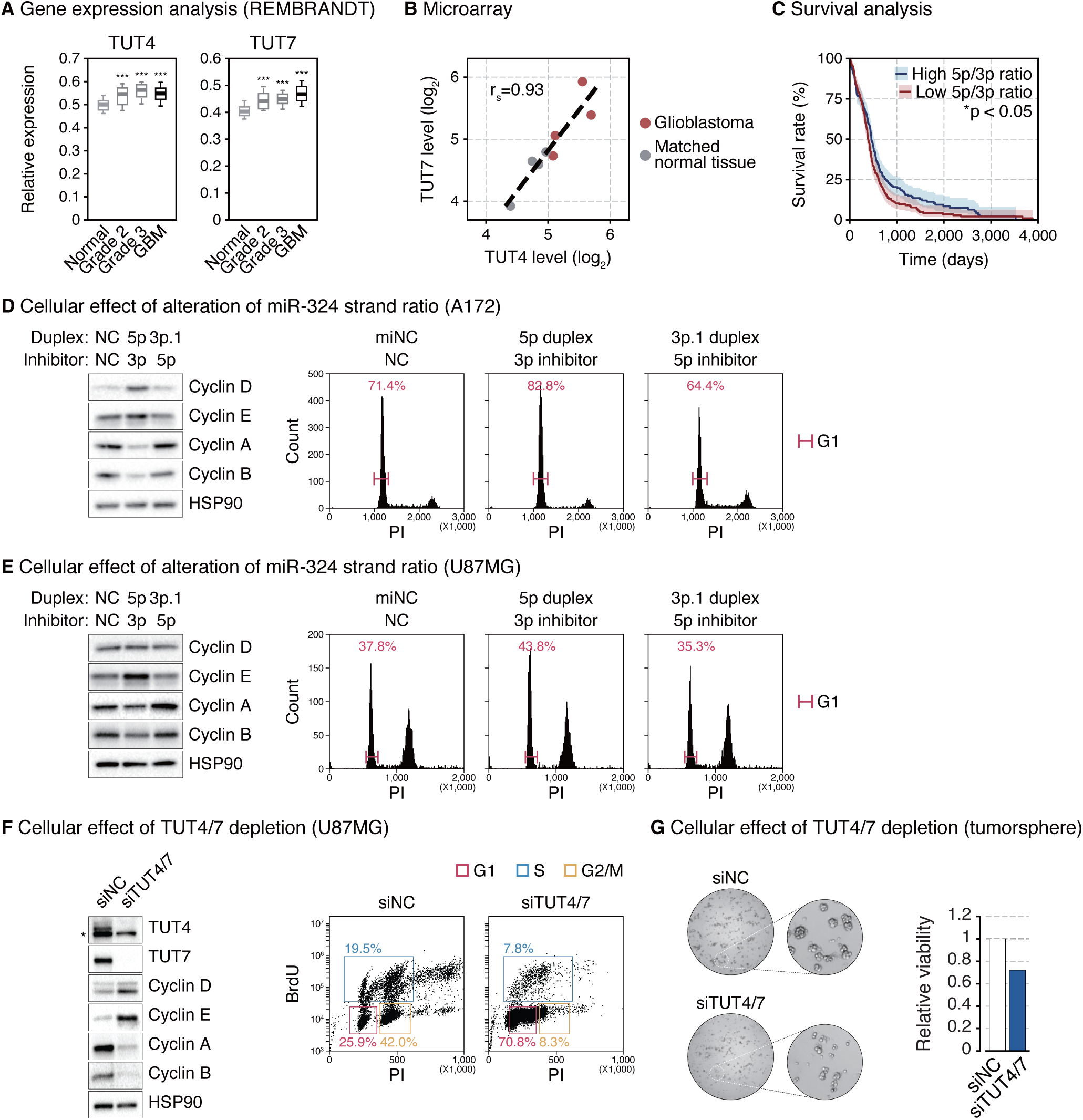
Perturbation of the miR-324 arm usage in glioblastoma suppresses cell cycle progression. **(A)** Expression levels of TUT4/7 in normal, lower grade glioma, and glioblastoma tissues from the REMBRANDT database. Normal, n = 28; grade 2, n = 65; grade 3, n = 58; glioblastoma, n = 228. ***p < 0.001 by one-way ANOVA with Tukey’s post hoc test for multiple comparisons. **(B)** Positive correlation between TUT4 and TUT7 expression levels detected in the microarray in glioblastoma and matched normal brain tissues. The linear regression is shown with a dashed line. r_s_, Spearman correlation coefficient. **(C)** The Kaplan-Meier survival curve stratified by high (n = 228) or low (n = 228) miR-324-5p/3p ratio of TCGA glioblastoma. Shades represent a 95% confidence interval. *p < 0.05 by the two-sided log-rank test. **(D–E)** Western blot of indicated cyclin proteins and cell cycle profile after disrupting the functional strand ratio of miR-324 in A172 (D) or in U87MG (E). **(F)** Western blot of indicated proteins and cell cycle after knocking down TUT4/7 in U87MG. *, a cross-reacting band. **(G)** Cell viability measured by the MTT assay after knocking down TUT4/7 in patient-derived glioblastoma tumorsphere (TS13-64). PI, propidium iodide. BrdU, bromodeoxyuridine.

**Figure S7.**
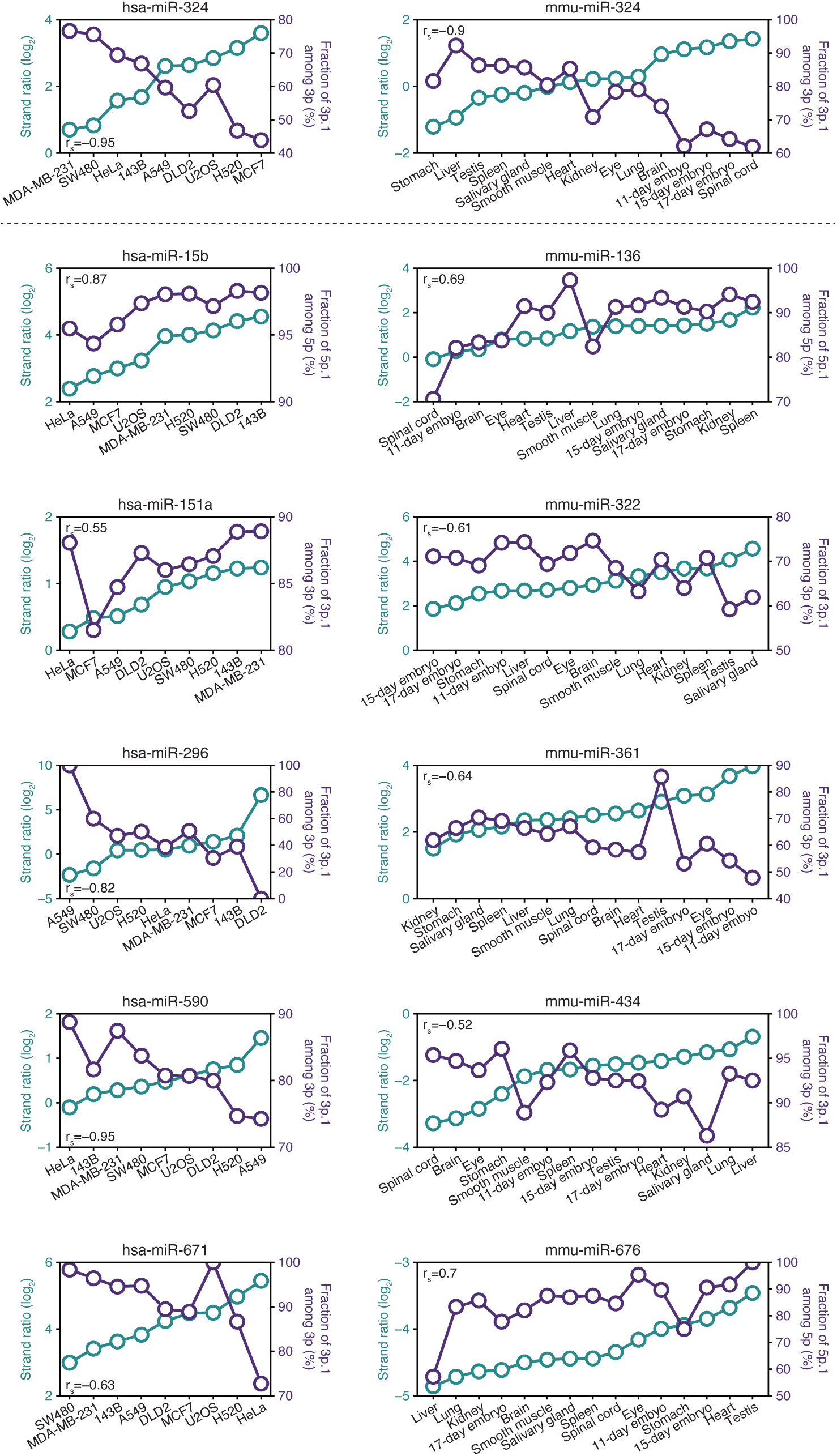
Strand ratios and 5′-isomiR fractions of many miRNAs change simultaneously depending on cell types. Scatter plots of the log_2_-transformed 5p/3p ratio of the most abundant 5′-isomiRs (cyan) and the fraction of the most abundant 5′-isomiRs among the indicated strand (purple).

